# De Novo Design of Allosteric Control into Rotary Motor V_1_-ATPase by Restoring Lost Function

**DOI:** 10.1101/2020.09.09.288571

**Authors:** Takahiro Kosugi, Tatsuya Iida, Mikio Tanabe, Ryota Iino, Nobuyasu Koga

## Abstract

Protein complexes exert various functions through allosterically controlled cooperative work. De novo design of allosteric control into protein complexes provides understanding of their working principles and potential tools for synthetic biology. Here, we hypothesized that an allosteric control can be created by restoring lost functions of pseudo-enzymes contained as subunits in protein complexes. This was demonstrated by computationally de novo designing ATP binding ability of the pseudo-enzyme subunits in a rotary molecular motor, V_1_-ATPase. Single molecule experiments with solved crystal structures revealed that the designed V_1_ is allosterically accelerated than the wild-type by the ATP binding to the created allosteric site and the rate is tunable by modulating the binding affinity. This work opened up an avenue for programming allosteric control into proteins exhibiting concerted functions.

---

Protein complexes exert their various functions through the cooperative work between their constituent subunits^1,2^. The orchestration between the subunits is enabled by the allosteric mechanism, in which a protein function in an active site is controlled by the response to stimuli that occurs at a site away from the active site^3^. The design of allosteric control into protein complexes to reveal their working principles and provide novel functionalities has been attempted^4–9^. One of these approaches is to create fusion proteins between the target protein, whose functions are to be controlled, and a protein undergoing conformational changes in response to stimuli, such as the binding of an effector molecule^6,7^ or light absorption^8,9^. Nakamura *et al*. created the remotely controlled linear motor protein with a light-sensitive domain^8^. In this study, we sought an approach to program allosteric control into protein complexes by creating binding sites for an allosteric effector molecule in the complexes.

We focused on pseudo-enzymes, which are homologs of some enzymes but are proven or predicted to have lost their enzymatic activity^10–12^. The overall structures of pseudo-enzymes are similar to those of the enzymes, but the conserved amino acids required for their functions are lost from the active sites; therefore, these sites are called pseudo-active sites. Interestingly, it has been reported that such pseudo-enzymes exhibit allosteric control when they form complexes ^10,11^. For example, a complex-forming pseudo-kinase—the pseudo-active site can bind ATP but has lost kinase activity—activates the catalytic function of a complex-forming partner protein, by ATP binding at the pseudo-active site^13,14^. Here, we hypothesized that an allostery can be de novo designed into protein complexes by restoring the lost function of pseudo-enzymes included in the complexes (Fig. 1a).

**Fig. 1.**
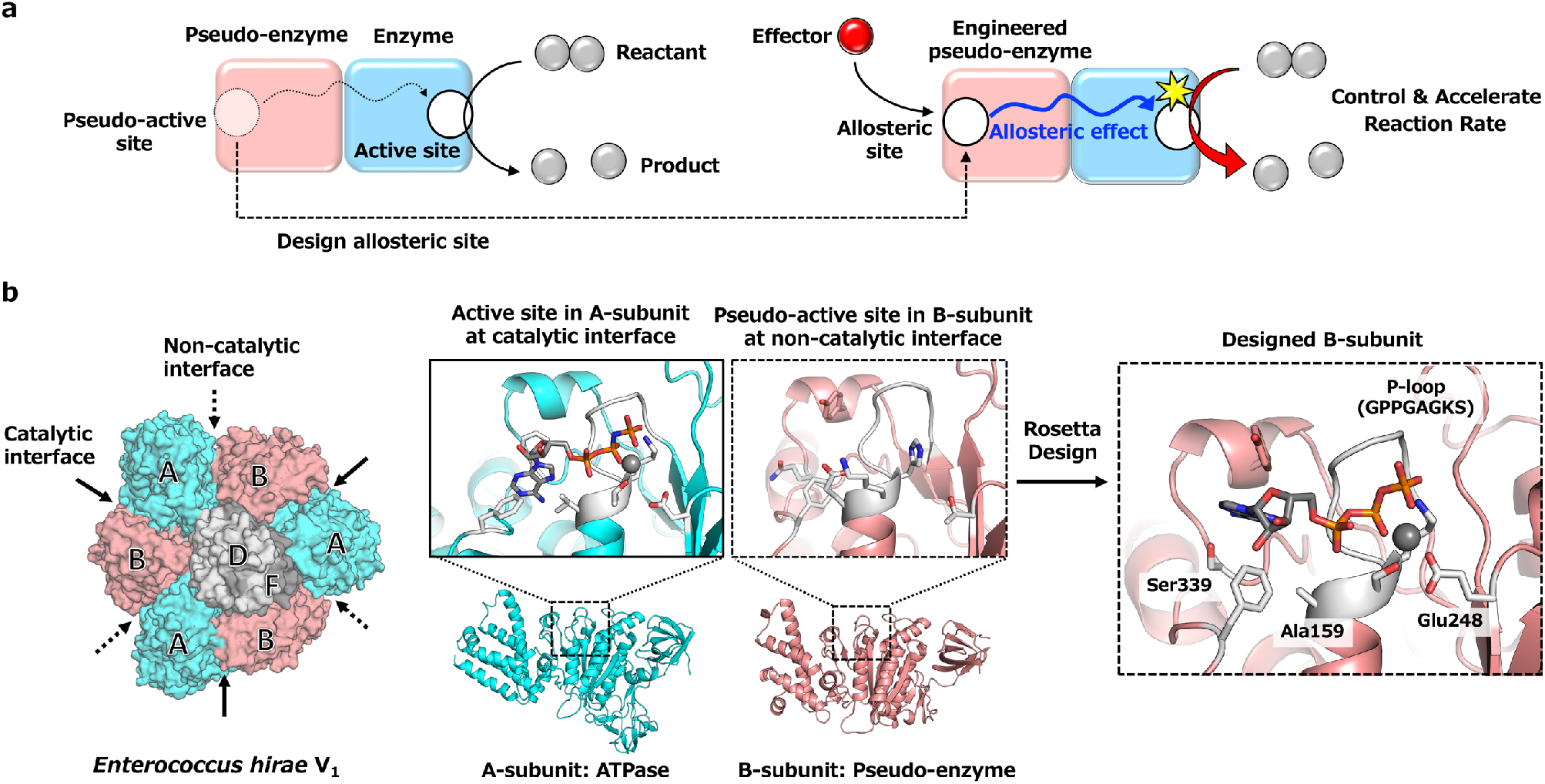
De Novo Design of an allosteric (ATP binding) site at the pseudo-active site of V_1_-ATPase. **a,** Strategy to de novo design of allosteric control into protein complexes by engineering the pseudo-active site. **b,** Overview of design scheme. Left: Catalytic and non-catalytic interfaces, indicated by solid and dotted arrows respectively, in the hexameric ring of V_1_ consisting of A-subunits (cyan) and their pseudo-enzyme B-subunits (salmon pink). The rotor of the D- and F- subunits (gray) is located in the center of the ring. Middle: The structures of A- and B-subunits with the active and pseudo-active sites, respectively. Right: An ATP binding site created at the pseudo-active site using the protein design software Rosetta; gray color residues were selected for the design (11 residue positions). The residues changed from the original sequence by the design are denoted with characters: the P-loop for binding to the phosphate group of ATP was built at the residue positions 151-158 with the amino acid sequence, GPPGAGKS; the Walker-B motif coordinating magnesium ion was built with glutamic acid at the residue position 248; the nucleotide-binding pocket was made with the alanine at 159, creating space for the binding of the sugar group of ATP (originally glutamic acid), and serine 339, making a hydrogen bond with the adenine ring.

A pseudo-enzyme is found in a protein complex, the rotary motor V_1_-ATPase (V_1_). V_1_ is a part of an ion pump V-ATPase which transports cations across the membrane by ATP hydrolysis-driven rotation^15^. The V_1_ consists of a rotor composed of the D- and F- subunits and the stator of the hexametric ring composed of three A-subunits and three B-subunits (Fig. 1b)^16^. The B-subunit is a pseudo-enzyme of the A-subunit and has homology with the A-subunit (e.g. for *Enterococcus hirae* V_1_-ATPase, the BLAST E-value between the subunits is 4 × 10^−22^) and the overall subunit structures resemble each other (Supplementary Fig. 1). However, the details of their sequences and structures are different, resulting in the two different interfaces between the A- and B- subunits in the A_3_B_3_ hexameric ring. One is the catalytic interface, at which the A-subunit has an ATP hydrolysis catalytic site, and the other is the non-catalytic interface, at which the B-subunit has a pseudo-active site, which does not hydrolyze or even bind ATP^16,17^(Supplementary Fig. 2).

The relation of the pseudo-active site to the activity at the active site is reported for V_1_, in which the mutations in the pseudo-active site decrease its activity^18,19^. Furthermore, a study on the rotary motor F_1_-ATPase from the thermophilic *Bacillus* PS3, which shares a common ancestor with V_1_-ATPase^20^, reported the relationship between the pseudo-active site and the active site in the rotational mechanism^21^. Similar to V_1_, F_1_ has the stator α_3_β_3_ ring complex, in which the β-subunit has ATP hydrolase ability, while the α-subunit, a pseudo-enzyme subunit of the β-subunit, can bind but not hydrolyze ATP^20,22^. The mutation in the pseudo-active site in the α-subunit, which significantly decrease the ATP binding ability, was found to cause F_1_ to have long pauses more frequently than the wild-type. This is likely because of the impeded release of ADP at the active site, indicating that the pseudo-active site away from the active site allosterically impacts on the active site’s function^21^. This study leads to the hypothesis that the pseudo-active site in the non-catalytic interface of V_1_, which does not have ATP binding ability^16,17^, can be a target for de novo design of a binding site for an allosteric effector molecule. We tested this hypothesis by computationally designing an ATP-binding site at the pseudo-active site in *Enterococcus hirae* V_1_-ATPase.

## Results

### Computational design of an allosteric site in the pseudo-active site

The pseudo-active site in the B-subunit does not have space for nucleotide binding or the well-known loop motif for phosphate binding, the Walker-A motif (GX_1_X_2_X_3_X_4_GK[T/S])^23,24^, also called a P-loop (Supplementary Fig. 2). Recently, computational methods for designing small molecule binding proteins have been developed, using Rosetta design software^25–27^. We attempted to computationally design an ATP binding site de novo together with a P-loop at the pseudo-active site in the B-subunit monomer, using Rosetta with a set of features for P-loops obtained from statistical analyses of naturally occurring proteins (Supplementary Fig. 3).

First, the backbone structure of the P-loop was built at the pseudo-active site by using the backbone of the A-subunit’s P-loop, considering the P-loop orientation feature (Supplementary Fig. 3a). Subsequently, side-chain conformations (amino acid sequences) of the P-loop and the surrounding residues, which have favorable interactions with ATP, are explored with various ATP conformations, the feature for native P-loops for the conserved amino acid (Gly) at X_3_ (Supplementary Fig. 3c), and the typical distances between the atoms of the P-loop and the phosphate atoms of ATP (Supplementary Fig. 3d). The resulting designed structures bound to an ATP were energetically minimized. This sequence design followed by energy minimization was iterated, and the designs with high ATP binding ability predicted by the Rosetta score were selected; the designs that lost the feature for the conserved backbone torsion pattern of native P-loops (Supplementary Fig. 3b) during the minimization step were abandoned. The ATP binding ability of 29 selected designs was further evaluated by short (10 ns) molecular dynamics simulations for the monomer (Supplementary Fig. 6). Finally, a resulting designed V_1_ was experimentally characterized. Details regarding the design procedure is described in Methods and Supplementary Figs. 4 and 5.

### Designed B-subunits forms a ring complex with A-subunits

The designed B-subunit (De), expressed with the A-subunit in *E. coli* using the plasmid pTR19-AB^28^ and purified by a Ni^2+^-affinity chromatography followed by size exclusion chromatography, formed a ring complex with the A-subunit (A_3_(De)_3_ ring complex) (Supplementary Fig. 7a). Subsequently, to evaluate the ATP binding ability of De, we introduced a double mutation in the A-subunit (K238A and T239A) to significantly impair ATP binding ability^29^. However, De did not form the A_3_(De)_3_ ring complex with the mutant A-subunit (Supplementary Fig. 7b). Therefore, we purified De as a monomer (Supplementary Fig. 7c) and the nucleotide binding ability was indirectly evaluated by thermal shift^30^ in circular dichroism spectroscopy in the presence or absence of nucleotides (Supplementary Fig. 8). The De monomer exhibited an increase of its melting temperature upon the addition of nucleotide, while the melting temperatures for the wild-type B-subunit monomer was almost the same in the presence and absence of nucleotides, suggesting that De has nucleotide binding ability. Thus, we attempted to determine the crystal structures of the A_3_(De)_3_ complex to prove the nucleotide-binding ability of De.

### The designed B-subunit binds to nucleotide

First, the A_3_(De)_3_ complex was crystallized in the absence of nucleotide and the structure was solved at 2.77 Å resolution, named A_3_(De)_3__empty. This showed the hexameric ring structure without nucleotide, which is the same as the wild-type structure (Fig. 2a). The structure of the designed site in the B-subunit in the crystal structure was almost identical to the computationally designed model (Supplementary Fig. 9a,b). When we incubated the nucleotide-free crystals with 20 μM AMP-PNP for 5 hours, we found an extra density corresponding to AMP-PNP in a catalytic site. The structure, A_3_(De)_3__(ANP)_1cat_, was solved at 3.44 Å resolution (Fig. 2b). Next, the nucleotide-free crystals were incubated for 5 hours with ADP by gradually increasing ADP concentration to 10 mM and the resulting structure, A_3_(De)_3__(ADP·Pi)_1cat_(ADP)_2cat,2non-cat_, was solved at 2.90 Å resolution. In A_3_(De)_3__(ADP·Pi)_1cat_(ADP)_2cat,2non-cat_, each of the three catalytic sites is occupied by ADP (one of the sites has ADP with a possible Pi) and each of the two sites out of the three design sites is with ADP (Fig. 2c). This crystal structure proved that the designed site in the non-catalytic interface has the nucleotide binding ability, although the binding mode was not same as we designed (Supplementary Fig. 9c,d). Furthermore, the nucleotide-free crystals were incubated overnight by gradually increasing the ADP concentration to 5 mM and two different structural states in an asymmetric unit were obtained from one dataset: one is A_3_(De)_3__(ADP)_3cat,1non-cat_, in which each of the three catalytic sites and one of the designed sites are occupied by ADP (Fig. 2d), and the other is A_3_(De)_3__(ADP)_3cat,2non-cat_, in which each of the three catalytic sites and each of the two designed sites are occupied by ADP (Fig. 2e). Although the resolution of these structures is relatively low (3.95 Å), the densities for the main chain Cα-trace and bound ADPs were clearly observed. The nucleotide-bound state of A_3_(De)_3__(ADP)_3cat,2non-cat_ is different from that of A_3_(De)_3__(ADP)_3cat,1non-cat_ in terms of nucleotide occupation at the designed site; these structures may provide the allosteric response upon ADP binding in the designed site (details are described later and in Fig. 5d).

**Fig. 2.**
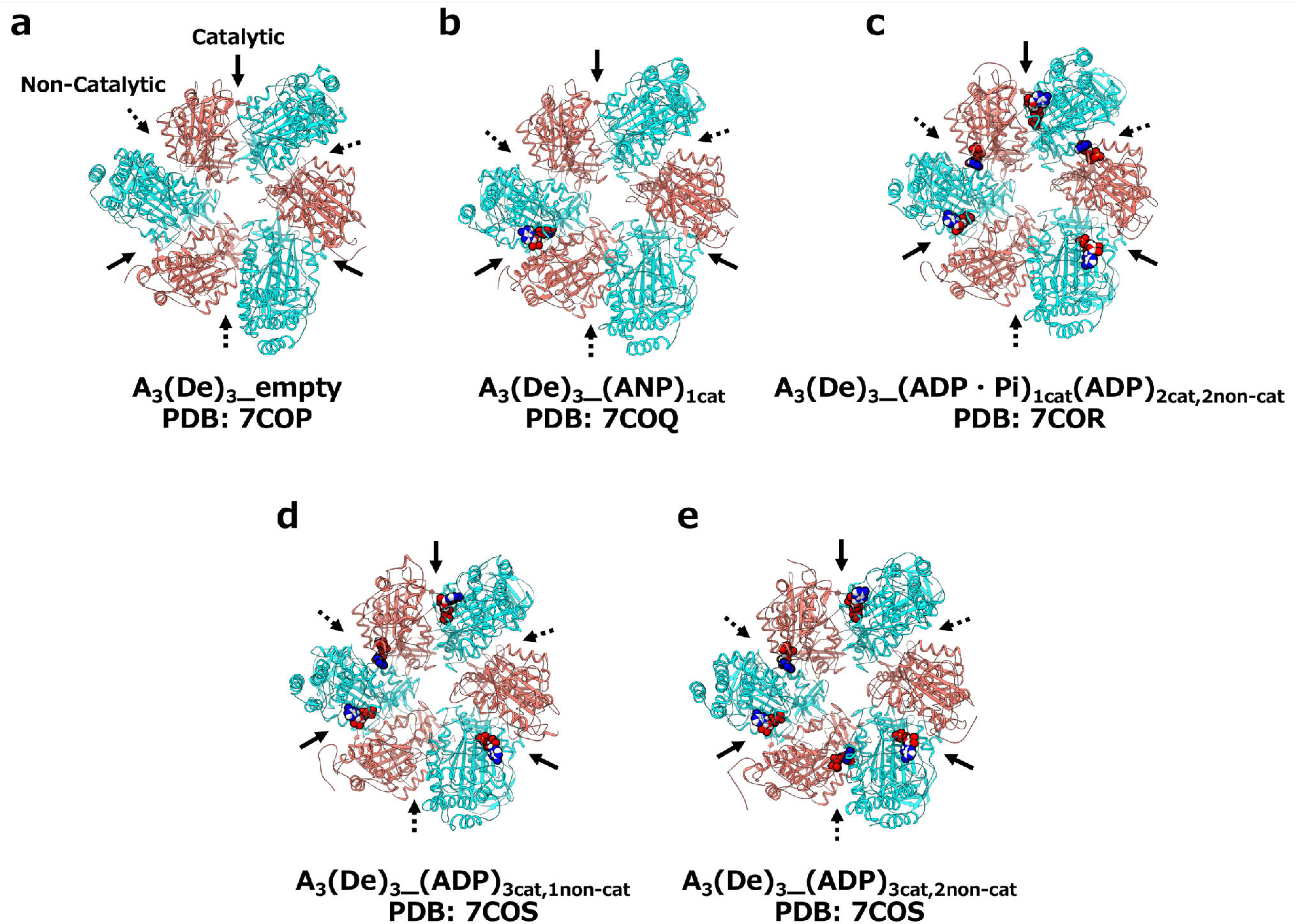
Solved five crystal structures of the A_3_(De)_3_ complex in various conditions. The presented structures are viewed from the C-terminal domain of the A- and designed B-subunits. **a,** A_3_(De)_3_ complex structure (De represents the design subunit) in the absence of nucleotides, solved at 2.77 Å resolution. This structure was named as A_3_(De)_3__empty. **b,** A_3_(De)_3_ complex structure bound to an AMP-PNP at a catalytic interface (3.44 Å resolution): A_3_(De)_3__ANP1cat. (C) A_3_(De)_3_ complex structure bound to 3 ADPs in the catalytic interfaces and 2 ADPs in the designed non-catalytic interfaces (2.9 Å resolution): A_3_(De)_3__(ADP·Pi)_1cat_(ADP)_2cat,2non-cat_. (D) A_3_(De)_3_ complex structure bound to 3 ADPs in the catalytic interfaces and an ADP in a designed non-catalytic interface (3.95 Å resolution): A_3_(De)_3__(ADP)_3cat,1non-cat_. (E) A_3_(De)_3_ complex structure bound to 3 ADPs in the catalytic interfaces and 2 ADPs in the designed non-catalytic interfaces (3.95 Å resolution): A_3_(De)_3__(ADP)_3cat,2non-cat_.

### Creation of ATP binding sites at the non-catalytic interfaces impacts on the catalytic interfaces

Fluorescence polarization measurements using the fluorescent-labeled AMP-PNP (Mant-AppNHp) and ADP (Mant-ADP), revealed that the nucleotide-binding affinities of the designed complex are much lower than those of the wild-type complex (Fig. 3a). The measured binding affinity of the A_3_B_3_ and A_3_(De)_3_ complexes respectively were 14.5 ± 5.7 nM and 2.03 ± 0.18 μM for AMP-PNP, and 64.2 ± 0.9 nM and 0.55 ± 0.003 μM for ADP. Note that the affinities of A_3_(De)_3_, measured in a range of relatively low nucleotide concentration, are expected for the first nucleotide binding to one of the catalytic interfaces, as supposed by the A_3_(De)_3__(ANP)_1cat_ structure, in which a single nucleotide was bound to one of the catalytic interfaces (Fig. 2b). Structure comparison of A_3_(De)_3__empty with the nucleotide-free wild-type A_3_B_3_ provides an interpretation for the decreased affinities (Fig. 3b). The P-loops in the three A-subunits in the wild-type A_3_B_3_ complex are classified into the two distinct conformations, bound and unbound forms (Supplementary Fig. 10). The P-loop in an A-subunit is the bound form and those in the other two A-subunits are the unbound form (Fig. 3b, top). The bound form is expected to have a higher binding affinity than the unbound form^16^. However, the P-loop conformations of all A-subunits in A_3_(De)_3__empty were found to be the unbound form, explaining the lower affinities of the catalytic interface. In other words, the creation of ATP binding sites in the non-catalytic interfaces changed the conformations of the ATP binding sites in the catalytic interfaces.

**Fig. 3.**
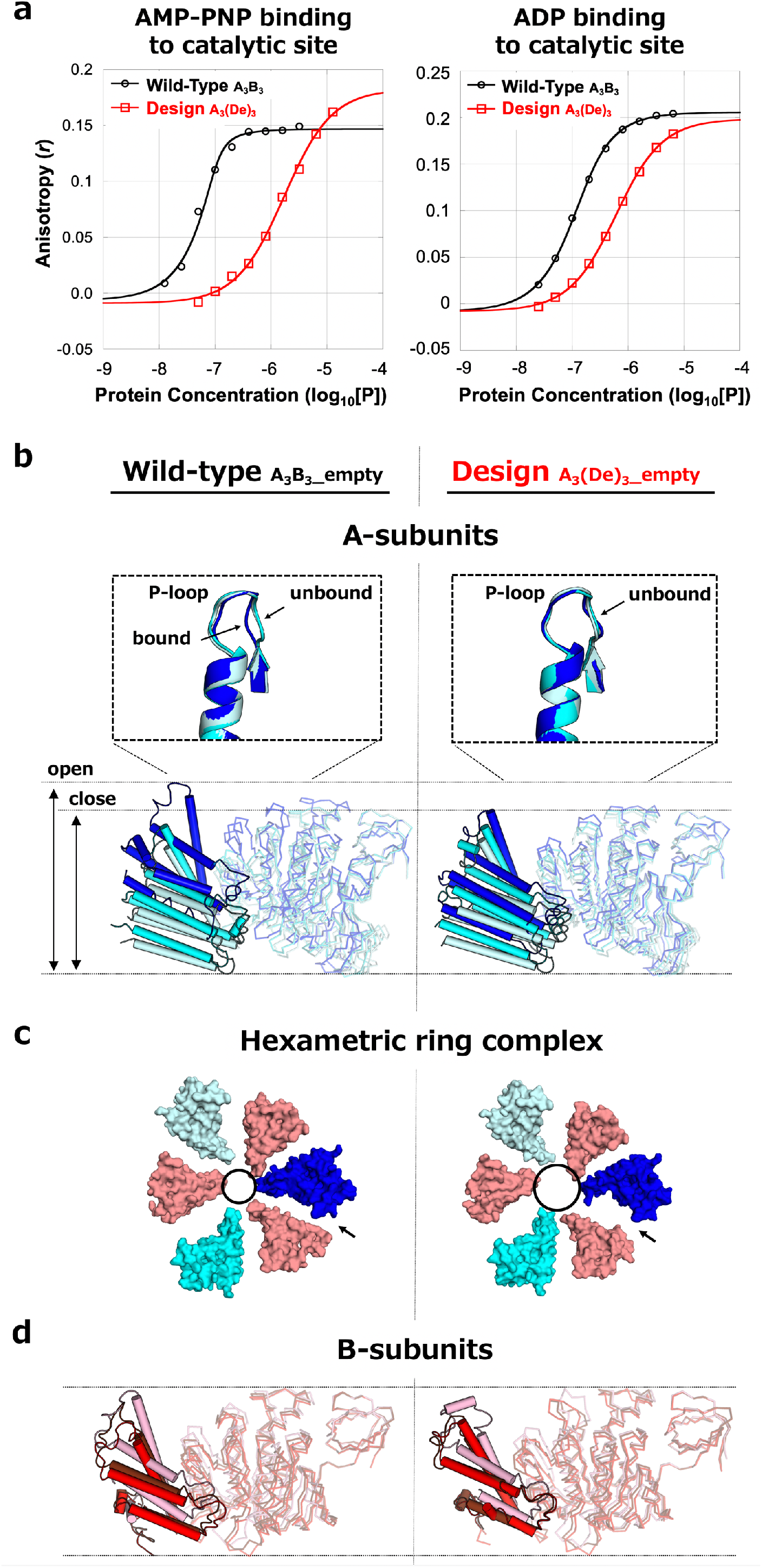
Creation of ATP binding sites in the B-subunit induced conformational changes of the A- subunit and the ring complex. **a,** The nucleotide binding affinities of the wild-type A_3_B_3_ and A_3_(De)_3_ complexes to AMP-PNP (top) and ADP (bottom), observed by fluorescent polarization experiments. **b,** Superpositions of the three A-subunit structures in the wild-type A_3_B_3_ (PDB: 3VR2) (left) and A_3_(De)_3__empty (PDB: 7COP) (right), using the β-barrel domains (residues 1-71), together with the close-up views for the structures around the P-loop. **c,** Ring complex conformations of the wild-type A_3_B_3_ complex (left) and A_3_(De)_3__empty (right). The C-terminal domains of A- and B- subunits viewed from the N-terminal β-barrel side, are shown. The circles and arrows show the central pore and the catalytic interface at which the largest conformational change is observed. **d,** Superpositions of the three B-subunit structures.

### Creation of ATP binding sites induces global conformational change of the A_3_(De)_3_ complex

The comparison of the A_3_(De)_3__empty with the nucleotide-free wild-type complex revealed that conformational changes occurred not only in the P-loop of the A-subunits but also in the overall structure of the A-subunits (Fig. 3b, bottom), although remarkable conformational changes were not found in any of the B-subunits (the structures of α-helical domains are locally different from the wild-type B-subunits due to their innate flexibility) (Fig. 3d). In the nucleotide-free wild-type complex, the A-subunit, of which the P-loop is the bound form, shows the closed conformation (the C-terminal domain bends toward the pore of the ring complex), and the other two A-subunits with the unbound form are in the open conformation^16^. However, all A-subunits in the design complex, A_3_(De)_3__empty, show the open conformation with the unbound form (Fig. 3b). Furthermore, these conformational changes broke the asymmetric arrangement of A- and B- subunits found in the wild-type ring complex^16^, resulting in a nearly symmetric arrangement with the expansion of the ring pore in the design complex (Fig. 3c). This observation leads to the hypothesis that the central rotor is difficult to retain in the expanded pore. As expected, the reconstitution experiments of the rotor (D- and F- subunits) and ring (A- and B- subunits) complex show that the reconstitution ratio was quite low (Supplementary Fig. 11). Interestingly, the conformational changes observed in A_3_(De)_3__empty were almost reverted by a nucleotide binding at a catalytic interface, as observed in A_3_(De)_3__(ANP)_1cat_ (Supplementary Fig. 11), and the complex reconstitution ratio was also recovered in the presence of nucleotide (Supplementary Fig. 12). The detailed comparisons of A_3_(De)_3__empty and A_3_(De)_3__(ANP)_1cat_ with the nucleotide-free wild-type complex structure^16^ are described in Supplementary Tables 1-4.

### The designed V_1_ rotates faster than the wild-type

Finally, we carried out single-molecule experiments to observe the rotation of the A_3_(De)_3_DF complex in various ATP concentrations ([ATP]s). The designed V_1_ was found to rotate unidirectionally in a counterclockwise fashion with discrete 120° steps, similar to the wild-type, but rotates faster than the wild-type at 100 μM ATP (Fig. 4a). Furthermore, in contrast to the wild-type, the rotation rate of the designed V_1_ exhibited an unique non-Michaelis-Menten type dependence on [ATP]. Rotation rates very similar to the wild-type were observed at the lowest (1 μM) and the highest (30 mM) [ATP], but the rotation was significantly accelerated in the range between the highest and lowest [ATP]s (Fig. 4b, top). At 100 μM ATP, the designed V_1_ showed the most accelerated rotation rate (115 ± 17 rps) compared with the wild-type (76 ± 4.8 rps). To the best of our knowledge, this is the first time that unidirectional movements of ATP-driven rotary molecular motors have been “overclocked” by protein engineering.

**Fig. 4.**
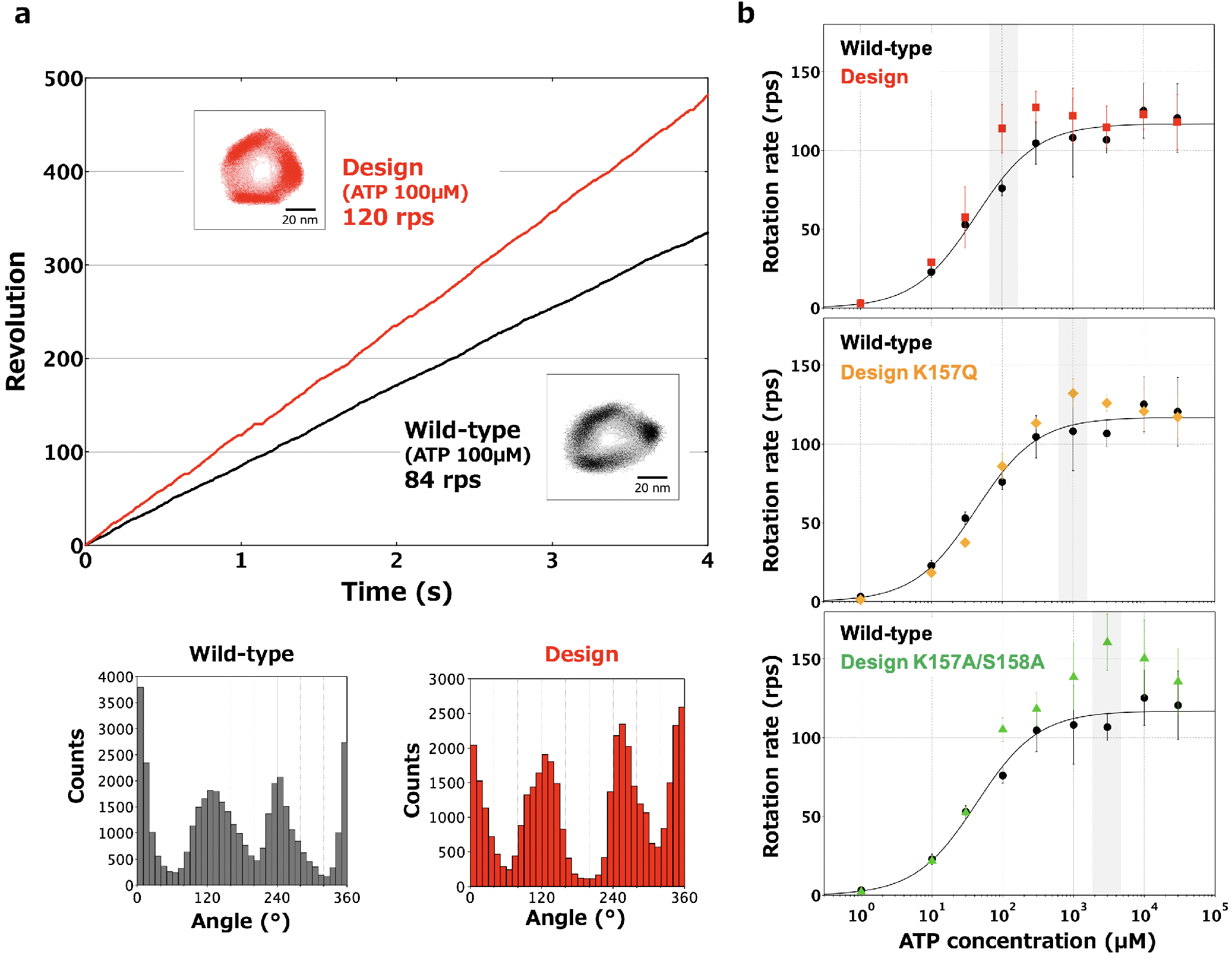
ATP binding to the designed site accelerates rotation rate allosterically. **a,** A typical rotation time course of the designed V_1_ (red) and that of the wild-type V_1_ (green)^28^, at 100 μM ATP. The insets show the rotation xy-trajectory. The angle distributions are shown at the bottom. **b,** [ATP] dependence of rotational rates for the wild-type (black)^28^, the designed V_1_ (red), the design mutant K157Q (orange) and the design double mutant K157A/S158A (green). The [ATP], at which the most accelerated rotation was observed, is highlighted in gray. The rates were plotted with averaged values using three molecules or more (Supplementary Table 5) and the error bars representing S. D.

Furthermore, we found that the [ATP], at which the designed V_1_ shows the maximal acceleration compared with the wild-type, can be tuned by modulating the nucleotide-binding affinity of the designed site (Fig 4b). The mutant K157Q at one of the conserved residues in P-loop motif in the designed site, expected to have a decreased binding affinity^31^, exhibited the non-Michaelis-Menten type rotation rate similar to the design, but notably, the [ATP], at which the most accelerated rotation was observed, shifted higher from that for the original design: 1 mM [ATP](Fig. 4b, middle). In addition, the double mutant K157A/S158A in the P-loop, which has a further decreased binding affinity (Note that the mutant is still expected to have a capability to bind ATP^29^), rotated in the similar fashion but the [ATP], at which the most accelerated rotation was observed, further shifted higher from that for the K157Q mutant: 3 mM [ATP] (Fig. 4b, bottom). Furthermore, the rotation rate at the ATP concentration was the highest (161 ± 18 rps) among those for the wild-type, the original design and the mutants (Fig. 4b). This observed correlation between the nucleotide binding affinity of the designed site and the [ATP] at which the most acceleration is observed strongly suggests the allosteric effect produced by the nucleotide-binding to the designed site.

### ADP-release at the catalytic site is facilitated allosterically

In the 120° step rotation, the designed V_1_ was found to have the main-pauses and sub-pauses before the 40° and 80° sub-steps respectively, which is the same as the wild-type V_1_^28^ (Fig. 5a). To reveal the mechanism of allosteric acceleration, we carried out dwell-time analyses of the two pauses at the high and low [ATP] (1 μM and 30 mM, respectively), in which the designed V_1_ rotated at a similar rate as the wild-type, and at the 100 μM [ATP], in which the design exhibited the most accelerated rotation. In the proposed rotation model for the wild-type^28^, the main-pause corresponds to the dwell-time waiting for ATP-binding, ATP-hydrolysis, and Pi-release, and the sub-pause corresponds to that for ADP-release. The main-pause time constants of the design at each measured [ATP] are roughly the same as those of the wild-type, irrespective of [ATP] (Fig. 5b). The sub-pause time constants for the wild-type stayed constant between 2.1~2.7 ms, at any [ATP]s, and the time constants for the design at the low and high [ATP] are similar to those for the wild-type. However, the time constant at the [ATP] (100 μM ATP), at which the designed V_1_ showed the most accelerated rate, significantly decreased to 1.0 ms (Fig. 5b). The double mutant (K157A/S158A) also exhibited behavior similar to the original design, and the sub-pause time constant was drastically decreased to 0.6 ms at 3 mM ATP. The rotation rates estimated from the measured time constants for the main- and sub-pauses, agreed with the observed rotation rates shown in Fig. 4b (Supplementary Table 6). All these results indicate that the origin of the acceleration is the facilitated ADP-release at the catalytic sites, which is generated through the allosteric effect triggered by nucleotides binding to the designed sites.

**Fig. 5.**
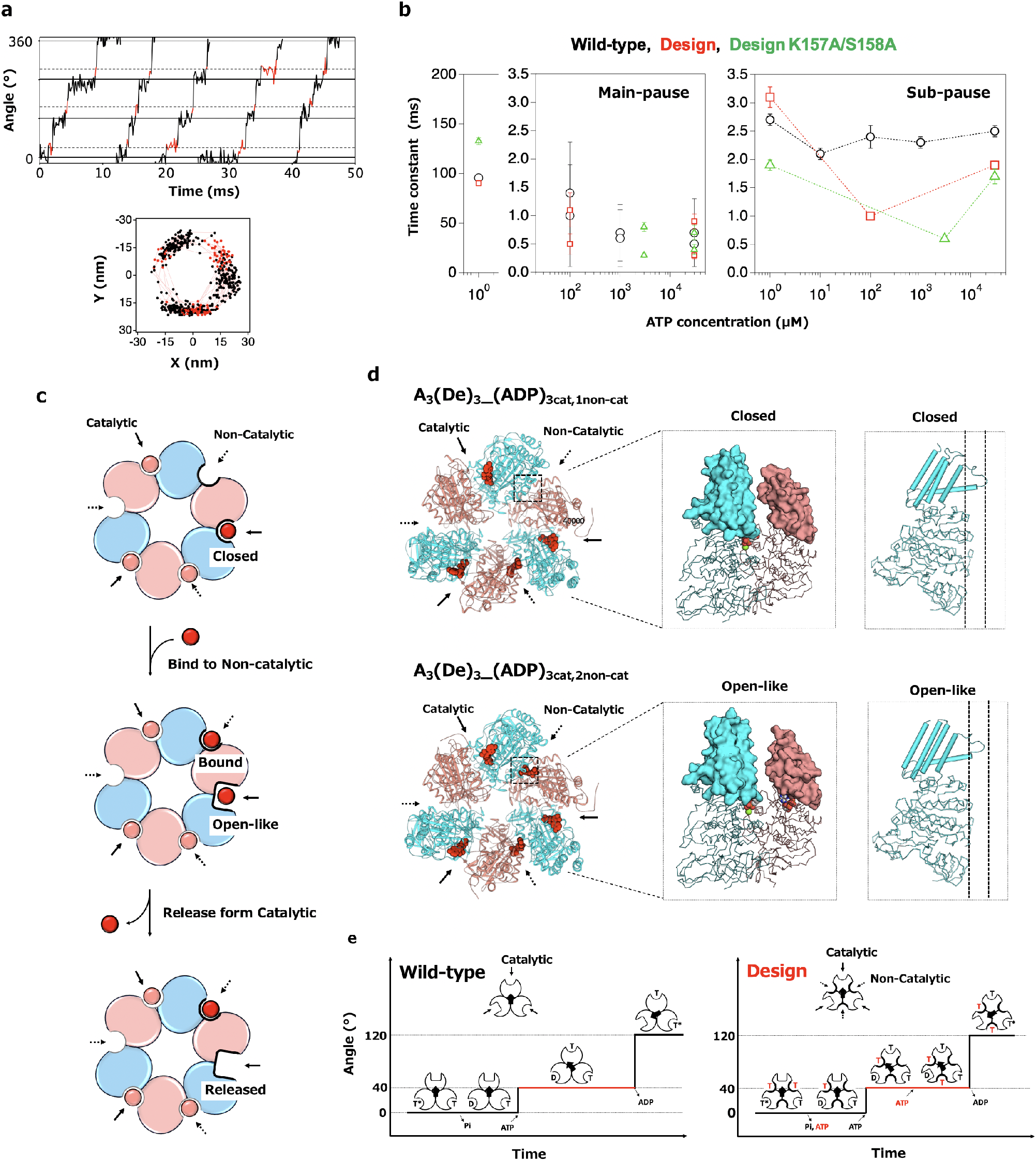
The mechanism of allosteric acceleration, revealed by analysis of rotation sub-steps and solved structures. **a,** A close-up rotation time course of the design at 100 μM ATP and the rotation xy-trajectory. The main-pause and sub-pause are black and red, respectively. **b,** Duration times at different [ATP] for the two pauses for the wild-type V_1_ (black), the designed V_1_ (red), and the design double mutant K157A/S158A (green). See Supplementary Fig. 13 for distributions of the duration time. Note that for the main pauses at 100-3000 μM ATP, two time constants were obtained for each [ATP] assuming consecutive reactions (see Supplementary Fig. 13). **c,** Structure-based interpretation on the ADP-release promoted by the allosteric effect. Ellipses indicate A-subunits (cyan) and designed B-subunits (salmon). Nucleotides are red (or salmon pink) circles. **d,** Comparison of A_3_(De)_3__(ADP)_3cat,1non-cat_ (top) and A_3_(De)_3__(ADP)_3cat,2non-cat_ (bottom). The hexameric structures viewed from the C-terminal domain of the A- and B-subunit (left) and the structures of the catalytic interfaces and the A-subunits, which form the closed and open-like conformations, respectively (middle and right). ADP molecules are shown as red spheres. **e,** A rotation scheme for the wild-type, proposed by Iida *et al*. ^28^(left). Proposed hypothetical rotation scheme for the design (right). ATP and ADP are represented by T and D, respectively. Red indicates nucleotides bound to non-catalytic interfaces.

Structural comparisons between the solved structures, A_3_(De)_3__(ADP)_3cat,1non-cat_ and A_3_(De)_3__(ADP)_3cat,2non-cat_, provide a structure-based interpretation for ADP-release promoted by the allosteric effect, although these structures are bound not with ATP but ADP, and do not contain the rotor. An ATP-binding at the designed site is suggested to induce conformational changes of the neighboring A-subunit and the catalytic interface from the closed conformation to the open-like conformation (Fig. 5c,d and Supplementary Tables 7 and 8), which creates space to facilitate ADP release.

### Possible model of the rotation scheme for designed V_1_

From the results described above, a rotation scheme for the designed V_1_ is proposed based on the scheme for the wild-type recently proposed by Iida *et al*. (Fig. 5e, left)^28^. For the wild-type, two or three catalytic sites are occupied at any time with ATP or its product(s) of hydrolysis. ATP-binding to an empty catalytic site triggers a 40° sub-step, and the subsequent release of ADP from the neighbor catalytic-site generates the 80° sub-step. The design may rotate in a similar scheme except for the ATP-binding to one or two designed sites at the non-catalytic interfaces, which facilitates ADP-release in the neighboring catalytic site through an allosteric effect (Fig. 5e, right). It is obvious that the allosteric effect does not emerge at low [ATP] since the designed sites are not able to bind ATP due to low affinity, but it is still puzzling why the allosteric effect is not observed in high [ATP]; the full nucleotide occupation in the three designed sites may suppress the allosteric effect, but, this should be verified in the future.

### Implications to native V_1_-ATPase and a common mechanism for rotary motors

The designed V_1_ not only exhibited the allosteric control over the rotation, but also provided possible designs and working principles for the native V_1_-ATPase. It is suggested that ancestral V_1_-ATPase existed as a homo hexameric ring, in which all subunits perform ATP binding and hydrolase functions, which has since evolved to form the current hetero hexameric ring containing the pseudo-enzyme subunit (B-subunit) that lost these functions^20^. Restoring ATP binding ability at the pseudo-active site may lead to an understanding of why the modern, descendant V_1_-ATPase lost its ATP binding and hydrolase functions. First, it is plausible that the non-catalytic interface observed in the modern V_1_-ATPase plays a role in attaining the Michaelis-Menten type rotation, in which the rotation rate is smoothly regulated along an [ATP] (Fig. 4b). The sudden increase and decrease of rotation rate at an [ATP] as observed in the designed V_1_ would not be preferable in terms of functional regulation by nature (this property can be beneficial for human since V_1_ can be engineered with the maximal rotation rate at an arbitrary [ATP]). Second, the non-catalytic interface may be a key factor for making the asymmetrical ring shape observed in the modern V_1_-ATPase in the absence of nucleotides, as we observed that the designed V_1_ forms a nearly symmetric ring structure (Fig. 3b,c). The asymmetrical ring shape is considered to be one of the key factors to realize the unidirectional rotation^16^ (it is reported that the N-terminal β barrel domains in the A- and B-subunits also play this role^32^). At the end, restoring ATP binding ability also implies a common mechanism for hexameric rotary motors of V_1_ and F_1_: the non-catalytic interface has the capability to provide allosteric control over ADP release at the neighboring catalytic interface, as observed in our design of V_1_ (Fig. 4b, and Fig. 5b,d) and the mutation of F_1_ described in the Introduction^21^.

### Conclusion

We succeeded in programing allosteric control into the molecular motor V_1_-ATPase by computational de novo design of a ATP binding site at the pseudo-active site in the non-catalytic interface of V_1_. Furthermore, the artificially designed V_1_ provided implications for design and working principles of the native V_1_-ATPase. Pseudo-enzymes are frequently found in native complex-forming proteins, e.g. F_1_-ATPase^22^, dynein^33^, kinesin^34^, 20S proteasome^35^, kinases^13^ and plant resistosome^36^. Engineering pseudo-active sites could be one of the promising approaches for de novo design of allosteric control into complex-forming proteins.

## Methods

### Computational design protocol

The B-subunit was computationally redesigned using the structure, chain E in PDB 3VR6. First, the pseudo P-loop in B-subunit (the residues 151-158) were replaced by the P-loop motif of A-subunit (the residues 232-239), by superimposing the A-subunit (chain B in the same PDB 3VR6) to the B-subunit with the orientation feature of P-loop shown in Supplementary Fig. 3a. Second, ATP binding modes were designed using Rosetta design software^37^ with Talaris2014 score function (Parameters for ATP-Mg^2+^ were determined using those for atom types already defined in Rosetta). In the design calculation, side-chain conformations for the residue positions of the P-loop and the surrounding residues (E169, T248, Q339, and F417), having favorable interactions with ATP, were explored with various ATP conformations generated by BCL software^38^ and with the distance constraints between the atoms of P-loop and the phosphate atoms of ATP (Supplementary Fig. 3d) (the amino acid at X_3_ in P-loop was fixed to Gly (Supplementary Fig. 3c)). Third, the designed B-subunit structures with ATP were minimized. The second and third steps were iterated 20 times and 800 different ATP binding modes were designed. Fourth, the designed structures that lost the feature for conserved backbone torsion pattern of P-loop (Supplementary Fig. 3b) were abandoned, and then those of which ATP binding score (Rosetta ddG score) are less than −8.0 were finally selected (29 designs).

### Molecular dynamics simulations

The 29 designs obtained by the computational design using Rosetta was further evaluated for their ATP binding ability by observing the stability of ATP in the designed site during short molecular dynamics (MD) simulations. The AMBER14 software suite^39^ was used for all MD simulations. The design models were used as the initial structures, of which hydrogen atoms were added by the LEaP module of AMBER14^39^. The simulation system contains a designed B-subunit monomer with ATP placed in a water box of approximately 82 Å × 112 Å × 100 Å. To neutralize the system, 15-17 sodium ions were put in the box. AMBER ff99SB sets and TIP3P were utilized for the protein and water molecules, respectively. Parameters for ATP molecule were adopted from a reference paper^40^. Long range electrostatic interactions were treated by the particle mesh Ewald (PME) method. Non-bonded interactions were cut off at 10 Å. After carrying out a short minimization to remove artificial repulsions in the initial structure, 10 ns MD simulations in a constant-NPT (300K, 1atm) ensemble were performed after the 100 ps heating stage with NVT ensemble (the time step is 2.0 fs and hydrogen atoms were constrained with SHAKE procedure). At the heating step, the temperature was raised gradually from 0 K to 300 K with the weak restraints (10 kcal/mol/A^2^) to the atoms of designed B-subunit. The MD simulation trajectory for each designed structure are shown in Supplementary Fig. 6 with the root mean square deviation (RMSD) values for the heavy atoms of ATP molecule from the minimized structure. Finally, a designed structure showing low RMSD value throughout the MD simulation was selected for experimental characterization.

### Expression and purification of the A_3_(De)_3_ Complex

A DNA fragment of the design was synthesized from the *ntpB* gene in pTR19-AB^28^ using megaprimer PCR method, and then the *ntpB* gene was replaced by this design fragment. The DNA sequence of design plasmid was confirmed by DNA sequencing analysis (Fasmac). *E. coli*. BL21* (DE3) competent cells were transformed with the plasmid and cultured at 30 °C for 20 hours in Super Broth (32 g/L Tryptone, 20 g/L yeast extract and 5 g/L sodium chloride) containing 100 μg/mL ampicillin and 2 mM isopropyl β-D-thiogalactopyranoside. Grown cells were spun down at 6,000 rpm for 10 minutes and washed twice with buffer A (20 mM potassium P_i_ (pH 7.0) and 100 mM NaCl). The cells were suspended in 15 mL of buffer A supplemented with 75 μL 100 mM phenylmethylsulfonyl fluoride (PMSF) solution and subsequently disrupted by sonication. After removing cell debris by centrifugation at 10,000 rpm for 20 minutes at 4 °C, the solution was filtered and applied to a Ni-NTA column. After washing with buffer B (20 mM potassium P_i_ (pH 7.0), 230 mM NaCl and 20 mM Imidazole), A_3_(De)_3_ complex was eluted with buffer C (20 mM potassium P_i_ (pH 7.0), 50 mM NaCl and 250mM Imidazole). The eluted fractions were concentrated with a Vivaspin20 5,000 MWCO (Sartorius) and then passed through a Superdex 200 Increase column (GE Helthcare) equilibrated with buffer D (20 mM MES-NaOH (pH 6.5), 100 mM KCl, 5 mM MgSO_4_, 0.1 mM DTT and 10% glycerol). The purified proteins were stored at −80 °C. The above described methods were also used for expression and purification of the wild-type A_3_B_3_ complex.

### Expression and purification of the design monomer

The designed B-subunit monomer was obtained by breaking the A_3_(De)_3_ complex sample in the presence of high concentration ATP. After expression and Ni-NTA purification of the A_3_(De)_3_ complex sample by the above described methods, the buffer of the A_3_(De)_3_ sample solution eluted from a Ni-NTA column was exchanged to buffer E (20 mM MES-NaOH (pH 6.5), 10% Glycerol, 100 mM KCl and 5 mM MgSO_4_) using PD-10 column (GE Helthcare). The sample solution mixed with 2 mM ATP was rocked for 30-40 minutes at 4 °C, filtered and applied to a Ni-NTA column. Because A-subunit has a His-tag and designed B-subunit does not, designed B-subunit can be selectively recovered in the flow through. In the flow through sample, Tris-HCl was added (100 mM final concentration; pH 8.5). The buffer of sample solution was exchanged to buffer F (20 mM Tris-HCl (pH 8.5), 10% Glycerol, 100 mM KCl and 5 mM MgSO_4_) by concentrating with a Vivaspin20 5,000 MWCO (Sartorius) and adding buffer F. The samples were passed through a Superdex 200 Increase column (GE Helthcare) equilibrated with buffer F. The purity of design monomer sample was confirmed by SDS-PAGE (Supplementary Fig. 7c).

### Expression and purification of the wild-type B-subunit

The wild-type B-subunit monomer was expressed with pTR19-B plasmid, which is constructed from the pTR19-AB plasmid^28^ by deleting the *ntpA* gene and adding His-tag to the *ntpB* gene, using the same protocol used for the A_3_(De)_3_ complex. The cells were suspended in 25 mL of buffer G (20 mM Tris-HCl (pH 8.5), 5% Glycerol, 0.7 M KCl, 5 mM MgSO_4_, 0.1 mM DTT and 20 mM Imidazole (pH8.5)) supplemented with 125 μL of 100 mM PMSF solution, and then disrupted by sonication. After removing cell debris by centrifugation at 10,000 rpm for 20 minutes at 4 °C, the solution was filtered and applied to a Ni-NTA column. After washing with buffer H (20 mM Tris-HCl (pH 8.5), 5% Glycerol, 0.7 M KCl, 5 mM MgSO_4_, 0.1 mM DTT and 20 mM Imidazole (pH8.5)), B-subunit monomer was eluted with buffer I (20 mM Tris-HCl (pH 8.5), 5% Glycerol, 0.7 M KCl, 5 mM MgSO_4_, 0.1 mM DTT and 250 mM Imidazole (pH8.5)). The eluted fractions were concentrated with a Vivaspin20 5,000 MWCO (Sartorius) and then passed through a Superdex 200 Increase column (GE Helthcare) equilibrated with buffer F. The purified proteins were stored at −80 °C.

### Expression and purification of the DF-subcomplex

The DF-subcomplex of *Enterococcus hirae* V_1_ were expressed in *E. coli*. BL21* (DE3) competent cells using the pTR19-D(M1G/T60C/R131C)F plasmid^28^. The transformed cells were cultured in Super Broth containing 100 μg/ml ampicillin at 37°C for 4-5 hours until OD_600_ reached 0.5, then the temperature was decreased to 30 °C and expression of DF-subcomplex was induced by the addition of 2 mM isopropyl β-D-thiogalactopyranoside. Cells were harvested 20 hours after induction by centrifugation at 6,000 rpm for 10 minutes. The cells were suspended in 20 mL of buffer J (20 mM potassium P_i_ (pH 8.0), 300 mM NaCl and 20 mM Imidazole) supplemented with 100 μL of 100 mM PMSF solution, and then disrupted by sonication. After removal of cell debris by centrifugation at 10,000 rpm for 20 minutes at 4 °C, the solution was filtered and applied to a Ni-NTA column. After washing with buffer J, DF-subcomplex was eluted with buffer K (20 mM potassium P_i_ (pH 8.0), 300 mM NaCl and 500 mM Imidazole). The eluted sample and TEV protease were mixed in 10:1 molar ratio and dialyzed against buffer L (20 mM potassium P_i_ (pH8.0), 50 mM NaCl, and 1 mM DTT) overnight. The dialyzed sample was spun down at 10,000 rpm for 20 minutes at 4 °C, and then applied to PD10 column for changing the buffer to buffer J. The eluted sample was applied to a Ni-NTA column and the flow thorough was collected. After adding 1 mM DTT, the sample was concentrated with a Vivaspin20 5,000 MWCO (Sartorius) and then passed through a Superdex 75 column (GE Helthcare) equilibrated with buffer M (20 mM Tris-HCl (pH 8.0), 150 mM NaCl). The purified proteins were stored at −80 °C. For single-molecule experiments, the cysteine residues introduced in D-subunit by the mutations T60C and R131C were biotinylated using the purified DF-subcomplex sample. The buffer of sample solution was changed to buffer N (20 mM potassium P_i_ (pH7.0), 150 mM NaCl) using PD10 column. The biotinylation regent (biotin-PEAC5-maleimide, Dojindo) was mixed into the purified DF-subcomplex sample solution with 3:1 molar ratio, and then incubated for 30 minutes at room temperature. Finally, DTT (10 mM final concentration) was added to the sample solution and the sample was stored at −80 °C. The purification results for gel filtration and SDS-PAGE are shown in Supplementary Fig. 7d.

### Expression and purification of the A_3_(De)_3_ for crystallization

The A_3_(De)_3_ protein sample for crystallization were prepared by cleaving the His-tag attached to the N-terminal of A-subunit. TEV protease cleavage site was inserted between the *ntpA* gene and His-tag in pTR19-AB^28^ by KOD-Plus-Mutagenesis Kit (TOYOBO). With this plasmid, the A_3_(De)_3_ sample was expressed and purified by using Ni-NTA column in the same protocol described above. The eluted sample and TEV protease were mixed in 10:1 molar ratio and dialyzed against buffer O (20 mM Tris-HCl (pH8.0) and 50 mM NaCl). This dialyzed samples were applied to a Ni-NTA column and the flow through was collected. Then, the sample was loaded onto a HiTrap Q HP column (GE Healthcare Life Sciences) equilibrated with buffer O, and then eluted with a linear gradient of buffer O with 50-1,000 mM NaCl in 20 min at flow rate of 1.0 ml min^−1^. The concentrated sample with a Vivaspin20 5,000 MWCO (Sartorius) was loaded onto a Superdex 200 Increase 10/300 GL column (GE Healthcare) equilibrated with buffer P (20 mM Tris-HCl (pH8.0), 150 mM NaCl and 2 mM DTT) at a flow rate of 0.5 ml min^−1^. The purified sample were concentrated with a Vivaspin500 5,000 MWCO.

### Preparation of the design mutants

The mutations were introduced by Quick Change Multi Site-Directed Mutagenesis Kit (Agilent Technologies). The purification and expression were carried out with the same method as the original design. The DNA sequence were confirmed by DNA sequencing analysis (Fasmac).

### Circular Dichroism measurement

Thermal denaturation experiments for the designed B-subunit and the wild-type B-subunit were carried out by using the Circular Dichroism spectrometer, J-1500KS (JASCO). Far-ultraviolet circular dichroism spectra at 220 nm along the increase of temperature in steps of 1.0 °C/min with 60 s of equilibration time were collected for 5 μM protein samples in buffer F in a 1-cm-path-length cuvette. The measurements were carried out in the absence of nucleotides or in the presence of 1 mM ADP, 1 mM ATP and 5 mM ATP after the incubation of the mixed solutions for 1 hour at 4 °C.

### Fluorescence polarization measurement for evaluating nucleotide-binding affinity of the A_3_(De)_3_ complex

Fluorescence polarization-based affinity measurements for the wild-type A_3_B_3_ complex and the designed A_3_(De)_3_ complex were performed using the fluorescent-labeled nucleotides, Mant-ADP and Mant-AppNHp (Jena Bioscience), in 100 nM. The changes in fluorescence anisotropy (*r*) of the fluorescent-labeled nucleotides mixed with the protein samples in Greiner black flat bottom 96 well plates, against the increase of the protein concentrations, were observed after 1 hour equilibration at room temperature on a Spark 10M (TECAN) using 360 nm excitation and 465 nm emission filters with 35 nm bandwidth filters. Buffer D was used for all measurement. Equilibrium dissociation constants (*K*_d_) were determined by the fitting to eq 1 with the anisotropy plots averaged over period of 10 min (20 measurements), where A is the experimentally measured anisotropy, A_f_ is anisotropy of the free ligand, A_b_ is the anisotropy of the fully bound ligand, [L]_T_ is the total ligand concentration, and [R]_T_ is the total protein concentration. The *K*_d_ values were determined by averaging the values from three independent measurements.

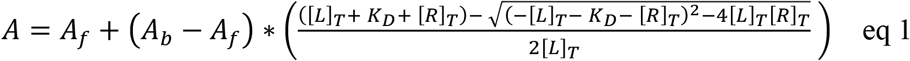

Note that the measured nucleotide affinities of the wild-type A_3_B_3_ complex are possibly underestimated by the binding at the second catalytic site, since the 100 nM nucleotide concentration, which is required to detect the fluorescence polarization of Mant, is not low enough against the binding affinity.

### Reconstitution experiments of the A_3_(De)_3_DF complex from the A_3_(De)_3_ complex and the DF-subcomplex

The A_3_(De)_3_DF complex was reconstituted from the A_3_(De)_3_ complex and the DF-subcomplex. The purified A_3_(De)_3_ and DF were mixed in a 1:5 molar ratio with the addition of MES-NaOH (pH6.0, 100 mM final concentration) and incubated for 2 hour at room temperature in the presence or absence of 20 μM AMP-PNP or ADP. The samples were filtered and passed through a Superdex 200 Increase column (GE Helthcare) equilibrated with buffer Q (20 mM MES-NaOH (pH 6.5), 10% Glycerol, 100 mM NaCl, 5 mM MgSO_4_ and 2 mM DTT). The reconstitution rate shown in Supplementary Fig. 11 was evaluated by SDS-PAGE for 1.5 μM, 14 μL of purified samples mixed with Tris-Glycine SDS buffer, and then quantified by ImageJ^41^ using eq 2, where A_WT_, A_Design_ are the optical densities of A-subunit in the wild-type complex or in the design complex, respectively, and D_WT_ and D_Design_ are those of D-subunit.

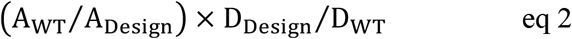

### Crystallization, data collection and structure determination

The sitting drop vapor diffusion method was used for crystallization. Crystals for A_3_(De)_3__empty were obtained by mixing 2.0 μL protein solution drop (10-15 mg/mL protein in buffer P) with 2.0 μL of reservoir solution (0.1 M Tris-HCl (pH 8.5), 20-24% PEG 3350 and 0.2 M Ammonium Acetate). The crystals were appeared in 1-2 weeks at 293K. The crystals were soaked in cryo-protectant solutions with an increasing concentration of 10% (v/v) glycerol. For A_3_(De)_3__(ANP)_1cat_, A_3_(De)_3__empty crystals were soaked for 5 hours in 20 μM AMP-PNP, 200 μM MgSO_4_ and 10% glycerol. For A_3_(De)_3__(ADP·Pi)_1cat_(ADP)_2cat,2non-cat_, A_3_(De)_3__empty crystals were soaked to ADP, MgCl_2_ and glycerol for 5 hours by gradually increasing the concentration to 10 mM, 10 mM and 10%, respectively. For A_3_(De)_3__(ADP)_3cat,1non-cat_ and A_3_(De)_3__(ADP)_3cat,2non-cat_, A_3_(De)_3__empty crystals were soaked to ADP, MgCl_2_ and glycerol overnight by gradually increasing the concentration to 5 mM, 5 mM and 10%, respectively.

The crystals were mounted on cryo-loops (Hampton Research), flash-cooled and stored in liquid nitrogen. All X-ray diffraction data were collected at the wavelength 1.1 Å on beamline BL-1A at Photon Factory (Tsukuba, Japan), from a single crystal at the cryogenic temperature (100K). The collected data were processed by using XDS^42^. The structure of A_3_(De)_3__empty and A_3_(De)_3_ with nucleotides were determined by molecular replacement method with Phaser^43^ using A_3_B_3_ complex from *Enterococcus hirae* (PDB 3VR2) and obtained A_3_(De)_3__empty structure as a search model, respectively. The initial model was iteratively refined with PHENIX^44^ and REFMAC5(CCP4 Suite)^45^ and manually corrected with COOT^46^. Figures are prepared by PyMOL^47^, CueMol2^48^ and Chimera^49^. The crystallographic and refinement statistics are summarized in Supplementary Table 9.

### Single-molecule experiments of the designed V_1_-ATPase

The protein sample was prepared by mixing the purified A_3_(De)_3_ and the biotinylated DF-subcomplex in a 1:5 molar ratio with the addition of MES-NaOH (pH6.0, 100mM final concentration), followed by the incubation in the presence of 200 μM ADP for 2 hours at room temperature. The sample was filtered and passed through a Superdex 200 Increase column (GE Helthcare) equilibrated with buffer Q and were concentrated to few μM with a Vivaspin500 5,000 MWCO. The samples were stored at −80 °C.

Single-molecule experiments were carried out by the method reported in the paper^28^. The flow cell was prepared by covering an untreated coverglass (18 × 18 mm^2^, Matsunami Glass) on a coverglass (24 × 32 mm^2^, Matsunami Glass) treated by overnight immersion in piranha solution (H_2_SO_4_/H_2_O_2_ = 3:1). After capturing the protein sample on the treated coverglass by His-tag, the streptavidin-coated 40-nm gold nanoparticle was attached to the biotinylated DF. The rotation of gold nanoparticle was observed by using an objective-type total internal reflection dark-field microscope^50^ constructed on an inverted microscope (IX-70, Olympus). The gold nanoparticles were illuminated by the evanescent field with the penetration depth of 100 nm from the glass surface. The scattered image of a rotating gold nanoparticle was recorded as a movie with a high-speed CMOS camera (FASTCAM 1024PCI, Photron) at 10,000 frames per second (fps) for almost all samples and at 27,000 fps for dwell time analyses of the double mutant at 3 mM ATP. During observation and recording under the microscope, ATP-regeneration system, in which ADP is rapidly regenerated to ATP by the coupling with dephosphorylation of phosphoenolpyruvate catalyzed by pyruvate kinase, was used to keep ATP concentration constant.

## Acknowledgement

We thank Prof. T. Murata at Chiba University for helpful suggestions; Dr. M. Kondo at ExCELLS and Dr. F. Kawai at Yamagata University for discussion about protein expression and purification; Dr. H. Ueno and Dr. Y. Minagawa at the University of Tokyo for advice about single-molecule experiments; Prof. T. Kinoshita at Osaka Prefecture University for discussion about pseudo-kinases; Prof. S. Akiyama at the Institute for Molecular Science and Prof. S. Takada at Kyoto University for comments on the manuscript; and Dr. R. Koga at ExCELLS for general discussion. We thank the Research Center for Computational Science (RCCS), Okazaki, Japan for providing its computational resources; the staff at the Photon Factory (PF) at KEK for assistance in synchrotron experiments and data collection for crystallographic analyses under the approval of PF program advisory committee (Proposal No. 2017G141); and for the support of the Basis for Supporting Innovative Drug Discovery and Life Science Research (BINDS) from AMED under Grant Number JP20am0101071 (supporting number BINDS0471). This work was supported by a Grant-in-Aid for Scientific Research on Innovative Areas “Molecular Engine” (JSPS KAKENHI Grant Number 18H05420 to T. K. and N. K. and 18H05424 to R. I.), by the National Institute for Natural Sciences (NINS), Okazaki Institute for Integrative Bioscience (Orion Project Grant Number 10341630611 to T.K. N. K. and R. I.), and by the Japan Science and Technology Agency (JST) Precursory Research for Embryonic Science and Technology (PRESTO, Grant Number JPMJPR13AD to N. K.).

## Author Contributions

T. K., R. I. and N. K. designed the research; T. K. computationally designed the ATP binding site; T. K. expressed and purified protein samples; T. K. performed biochemical measurements; T. K. and M. T. performed crystallography experiments and analyzed the data; T. K. and T. I. performed single-molecule experiments and analyzed the data; T. K. and N. K. wrote the manuscript; and T. K. coordinated the overall research. All authors discussed the results and commented on the manuscript.

## Additional Information

**Supplementary Information** is available for this paper.

