## Supplementary Information for "De Novo Design of Allosteric Control into Rotary Motor V_1_-ATPase by Restoring Lost Function"

#### Supplementary Figures

1. Structural similarities between the A-subunit and its pseudo-enzyme, B-subunit
2. Structural differences between the active site in the A-subunit and the pseudo-active site in the B-subunit
3. Conserved features of P-loop motifs and typical binding atom-pair distances with phosphate group of ATP, used for designing ATP-binding site in the B-subunit's pseudo-active site
4. Flowchart for designing ATP binding site in the B-subunit's pseudo-active site
5. Design target residues to create ATP binding site in the B-subunit's pseudo active site
6. ATP binding abilities of the 29 designs, evaluated by short MD simulations
7. Gel filtration and SDS-PAGE for the $A_3(De)_3$ complex, complex of the A-subunit double mutant K238A/T239A and the designed subunit, the design monomer and the DF-subcomplex
8. Thermal shifts of the wild-type B-subunit and the designed monomers by nucleotide binding
9. Structural comparisons of the computational design model with the solved crystal structures
10. Bound and unbound forms observed in P-loop motifs in the wild-type structure (PDB: 3VR3)
11. Design of ATP binding sites in the B-subunit induced conformational changes of the ring complex
12. Complex formation ability of the $A_3B_3$ and $A_3(De)_3$ hexametric ring with the central axis (DF-subcomplex) in the presence or absence of nucleotide
13. Duration time distributions with calculated time constants for main- and sub-pauses for the designed $V_1$ and the design double mutant K157A/S158A

#### Supplementary Tables

1. Structural comparison of A-subunits between the nucleotide-free $A_3B_3$ complex of the wild-type (3VR2) and the designed $V_1$ ($A_3(De)_3$_empty).
2. Structural comparison of catalytic interfaces between the nucleotide-free $A_3B_3$ complex of the wild-type (3VR2) and the designed $V_1$ ($A_3(De)_3$_empty)
3. Structural comparison of the A-subunits between the nucleotide-free $A_3B_3$ complex of the wild-type (3VR2) and the designed $V_1$ ($A_3(De)_3$_$(ANP)_{1cat}$)
4. Structural comparison of the catalytic interfaces between the nucleotide-free $A_3B_3$ complex of the wild-type (3VR2) and the designed $V_1$ ($A_3(De)_3$_$(ANP)_{1cat}$)
5. Number of molecules used for measurement of average rotation rates in the single-molecule experiments
6. Comparison of rotation rates estimated from the slope in the time course of rotation and those estimated from the dwell time constants for the main- and sub-pauses
7. Structural comparison of the A-subunits between the nucleotide-free $A_3B_3$ complex of the wild-type (3VR2) and the designed $V_1$ ($A_3(De)_3$_$(ADP)_{3cat,1non\text{-}cat}$ and $A_3(De)_3$_$(ADP)_{3cat,2non\text{-}cat}$)
8. Structural comparison of the catalytic interfaces between the nucleotide-free $A_3B_3$ complex of the wild-type (3VR2) and the designed $V_1$ ($A_3(De)_3$_$(ADP)_{3cat,1non\text{-}cat}$ and $A_3(De)_3$_$(ADP)_{3cat,2non\text{-}cat}$)
9. Data collection and refinement statistics of crystal structures

#### Supplementary Method

1. Rosetta Scripts XML file for design of ATP binding site

#### Supplementary References

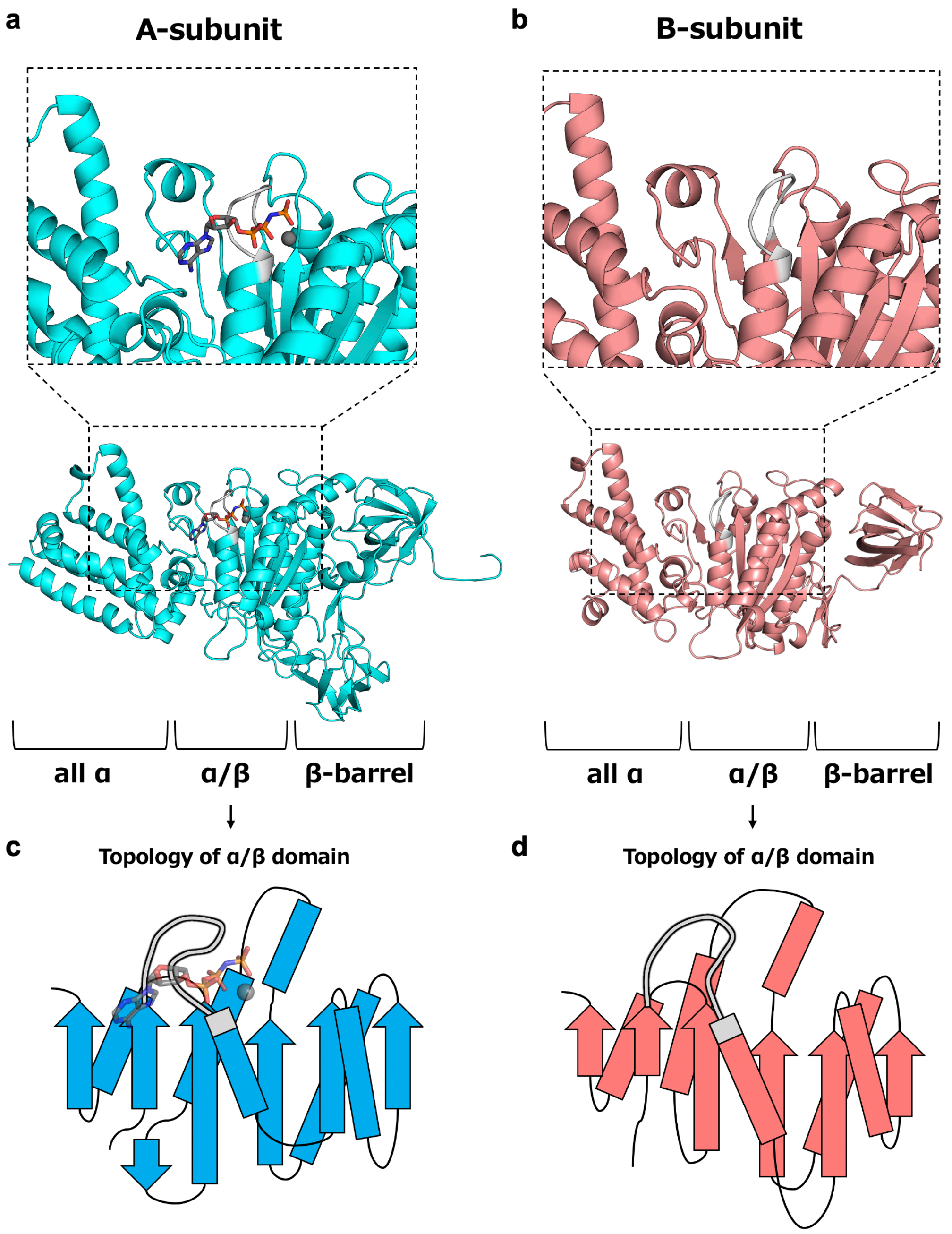


**Supplementary Fig. 1: Structural similarities between the A-subunit and its pseudo-enzyme, B-subunit.** The A- and B-subunit are presented at left and right, respectively. **a,b**, Entire structures of the A- and B- subunits with the close-up views of structures around the active site in the A-subunit and the pseudo-active site in the B-subunit. Both subunits consist of all-α-, α/β-, and β-barrel domains and have similar backbone structure (TM-score evaluated by MICAN[1] is 0.87). The spatial arrangement of secondary structures around the pseudo-active site in the B-subunit is nearly the same as those of the active site in the A-subunit. **c,d**, Topology schematics of α/β domains of the A- and B- subunits. The α/β domain of the B-subunit (residue 78-362) has an almost identical topology with that of the A-subunit (residue 72-449).

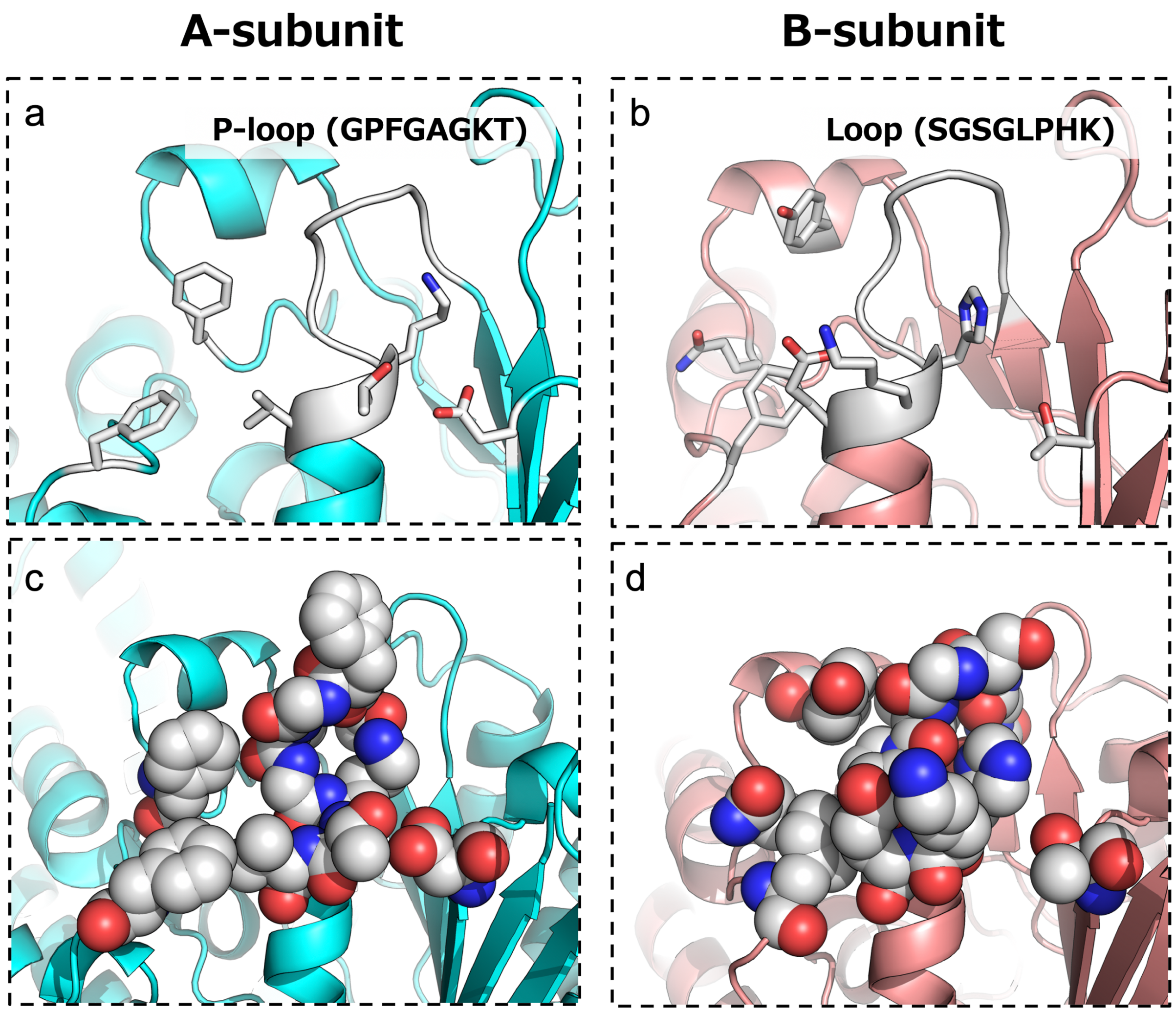


**Supplementary Fig. 2: Structural differences between the active site in the A-subunit and the pseudo-active site in the B-subunit.** The A- and B-subunit are presented at left and right, respectively. The residues for nucleotide-binding in the A-subunit's active site are shown by stick in **a** and sphere in **c**. The corresponding residues in the B-subunit's pseudo-active site are shown by the same representations in **b** and **d**. The A-subunit's active site has a P-loop motif (GXXXXGKT/S) for binding to phosphates of ATP (See **a**), and has a space for binding to the sugar and base of ATP molecule (See **c**). In contrast, these features are not observed in the B-subunit's pseudo-active site: the loop corresponding to the P-loop does not have the conserved sequence (See **b**) and the typical backbone geometries (See Supplementary Fig. 3a,b); and there is no space for ATP binding (See **d**).

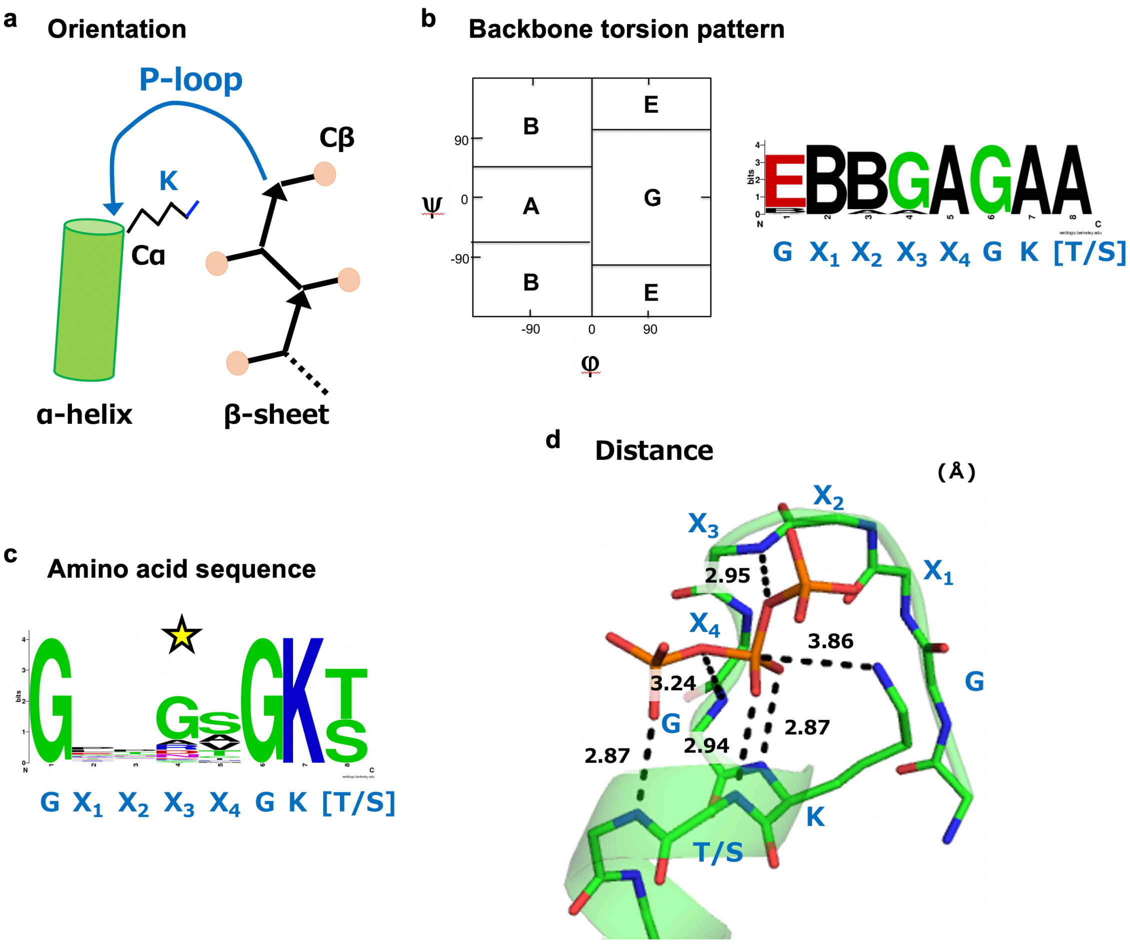


**Supplementary Fig. 3: Conserved features of P-loop motifs and typical binding atom-pair distances with phosphate group of ATP, used for designing ATP-binding site in the B-subunit's pseudo-active site.** P-loop motifs are found in the connection of successive secondary structure elements from β-strand to α-helix. Statistical analysis of 52 X-ray structures containing P-loop motif ($GX_1X_2X_3X_4G$[T/S]), collected from the PISCES server[2] with resolution ≦ 2.0 Å, R-factor ≦ 0.25, and sequence identity ≦ 25 %, revealed the three conserved features and the typical distance with the phosphate group of ATP. **a,** The orientation. The $C_\beta$ atom of the last strand residue immediately before the P-loop locates at the opposite side to that at which the $C_\alpha$ atom of the conserved Lys locates. **b,** The backbone torsion pattern. The residues in P-loop have the backbone torsion pattern, represented by ABEGO[3] torsion bins: EBBGAGAA (as shown in left, the torsion bins A and B are the α-helix and β-sheet regions; G and E are the positive phi regions; and O is the cis peptide bond). **c,** The amino acid sequence. P-loop motif is defined by the conserved sequence, $GX_1X_2X_3X_4$GKT/S, but the motif also has conserved residues in the region from $X_1$ to $X_4$. Especially $X_3$ indicated by star is Gly. **d,** The distances between the atoms of P-loop and the phosphate atoms of ATP. Each of distances was obtained by averaging the corresponding distances of P-loop motifs found in nature.

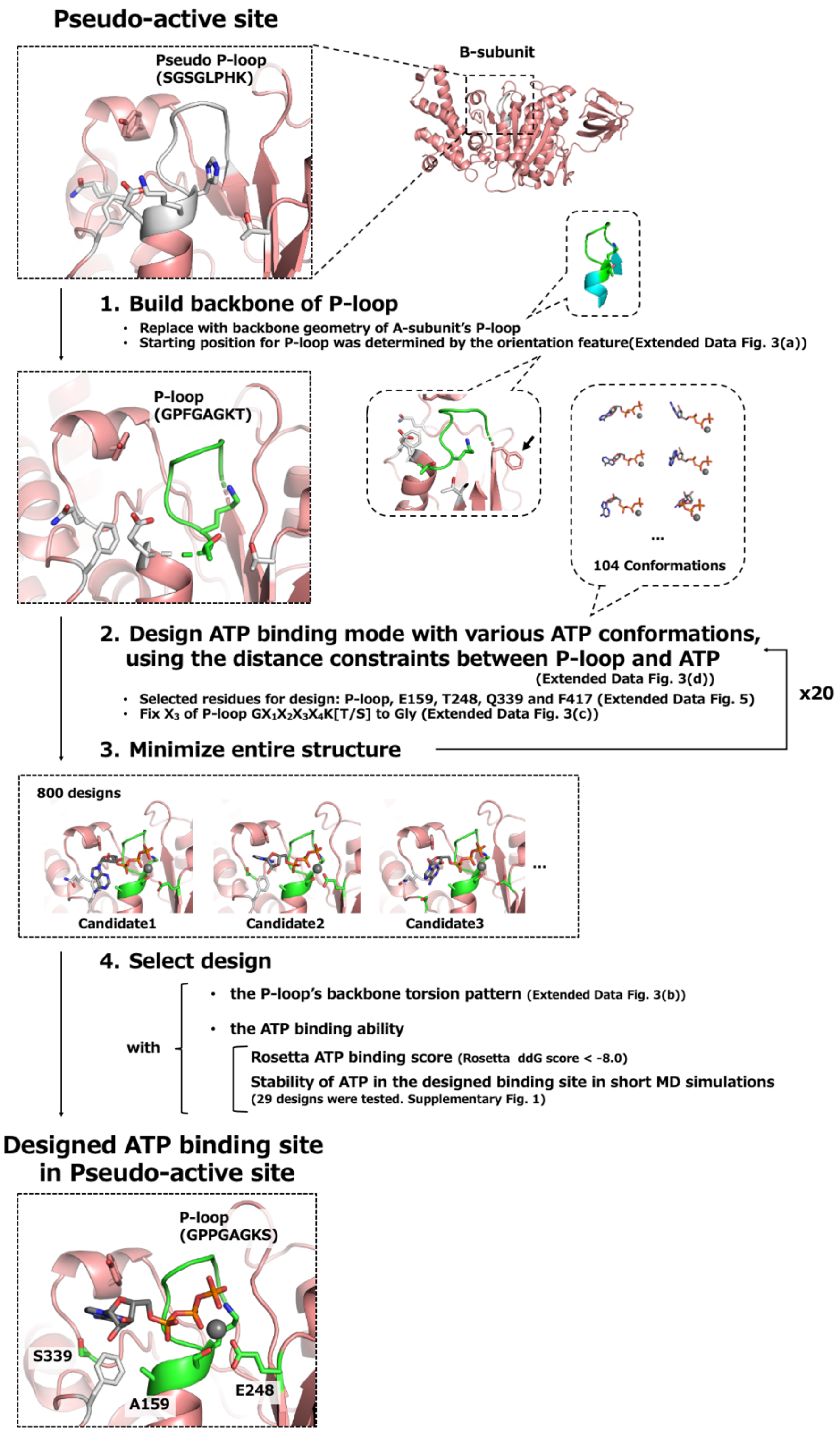


**Supplementary Fig. 4: Flowchart for designing ATP binding site in the B-subunit's pseudo-active site.**

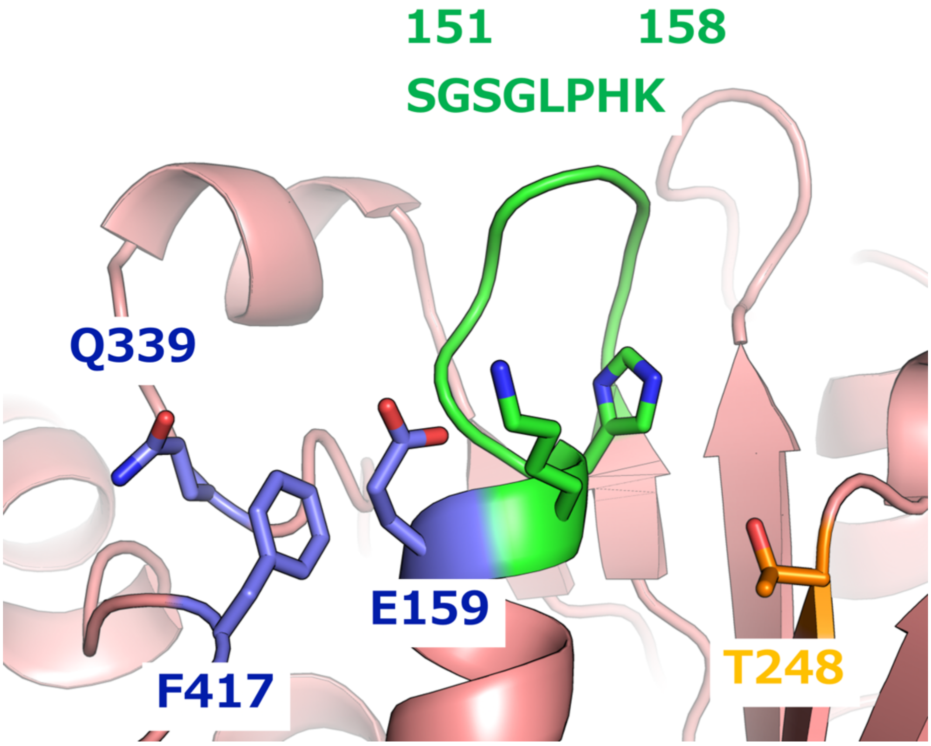


**Supplementary Fig. 5: Design target residues to create ATP binding site in the B-subunit's pseudo active site.** Residue positions in the pseudo P-loop from 151 to 158 (green) were designed as a P-loop motif (G$X_1X_2X_3X_4$G[T/S]). The amino acid type at $X_3$ position in the P-loop motif was also fixed to glycine, because of the high conservation (Supplementary Fig. 3c). T248 (orange) is the position for the Walker B motif, this position was designed using aspartic or glutamic acid residue. For making a space for nucleotide binding, the three residues (purple) were selected as minimal as possible because the redesigned $V_1$ lost the ability to form the complex when more residues were selected.

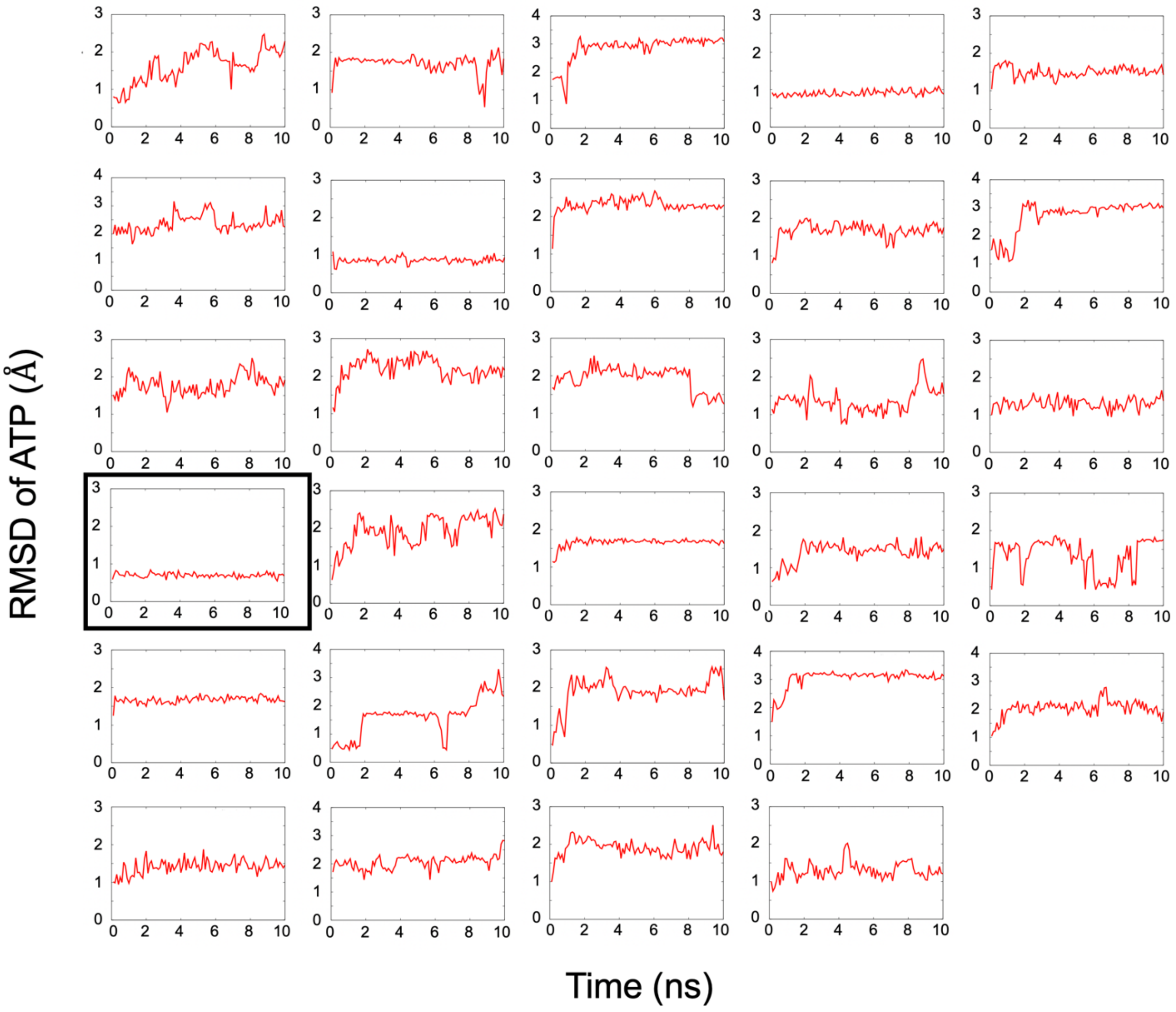


**Supplementary Fig. 6: ATP binding abilities of the 29 designs, evaluated by short MD simulations.** In each panel, RMSD values calculated using heavy atoms of ATP molecule in a designed structure are plotted along the time course. The MD trajectory surrounded by the thick line frame is the design selected for experimental characterizations.

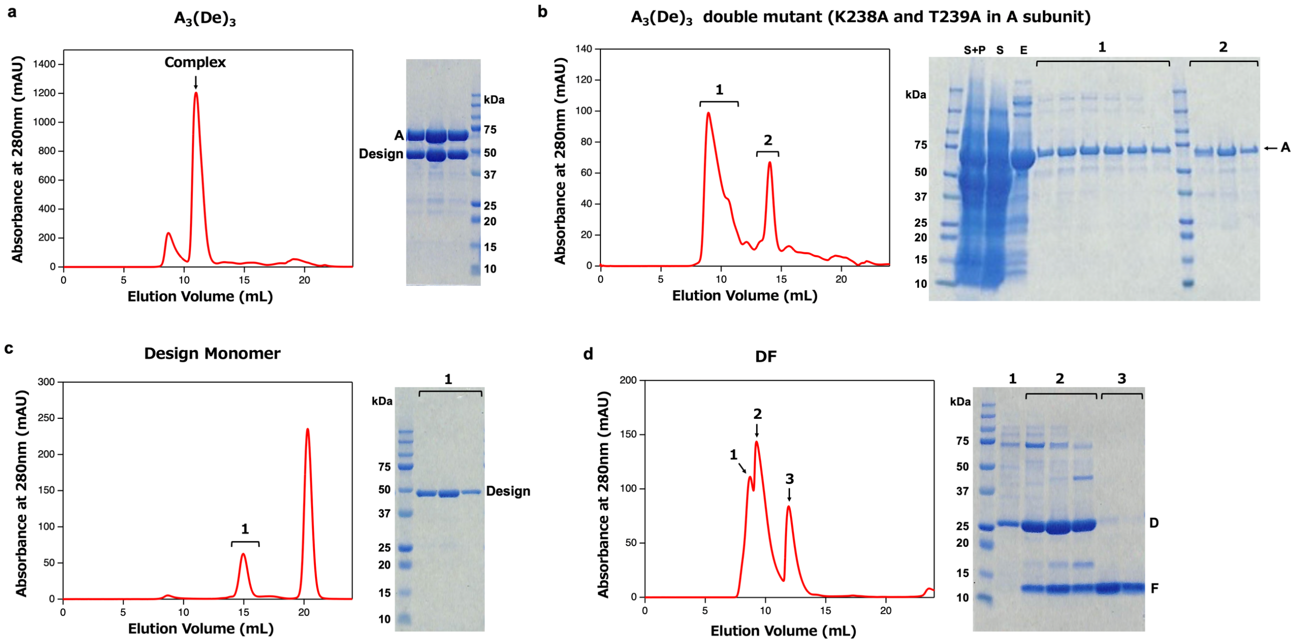


**Supplementary Fig. 7: Gel filtration and SDS-PAGE for the $A_3(De)_3$ complex, complex of the A-subunit double mutant K238A/T239A and the designed subunit, the design monomer and the DF-subcomplex.** In the figure of SDS-PAGE, S+P, S and E indicate supernatant + pellet, supernatant and elution, respectively. **a,** $A_3(De)_3$ complex; the $A_3(De)_3$ was purified as the complex state. **b,** Complex of the A-subunit double mutant K238A/T239A and the design; the A-subunit double mutant K238A/T239A does not form the complex with our designed subunit. In SDS-PAGE, S+P, S and E indicate supernatant + pellet, supernatant and elution, respectively. **c,** Design monomer; Purification of the design monomer. The $A_3(De)_3$ complex is broken by adding excessive ATP and design monomer was purified. The design monomer was collected from the peak of gel filtration (left) and the purity (only design monomer without the A-subunit) was evaluated by SDS-PAGE (right). **d,** DF-subcomplex; Purification of the DF subcomplex. The DF-subcomplex was purified as a complex.

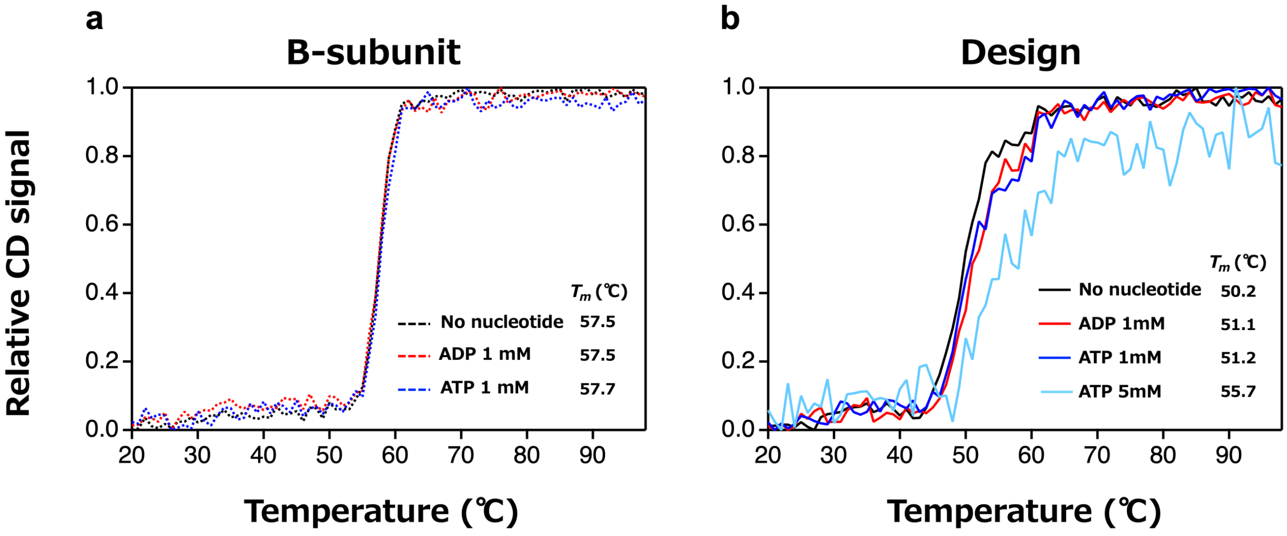


**Supplementary Fig. 8: Thermal shifts of the wild-type B-subunit and the designed monomers by nucleotide binding.** Thermal denaturation curves for the wild-type B-subunit (**a**) and the design monomer (**b**) in the presence and absence of nucleotide, measured by Circular Dichroism (CD) at 220nm, are shown. The CD signal values in the denaturation curves were normalized between 0 and 1 by min-max normalization: (CD signal – lowest CD signal)/(highest CD signal – lowest CD signal)). While the wild-type B-subunit shows almost the same melting temperatures both in the presence and absence of nucleotide, the design monomer exhibits higher melting temperatures in the presence of ATP and ADP than those in the absence of nucleotide. The melting temperature $T_m$ values for the B-subunit and the designed monomer were obtained from the denaturation curves by non-linear least-squares analysis using a two-state unfolding and linear extrapolation model.

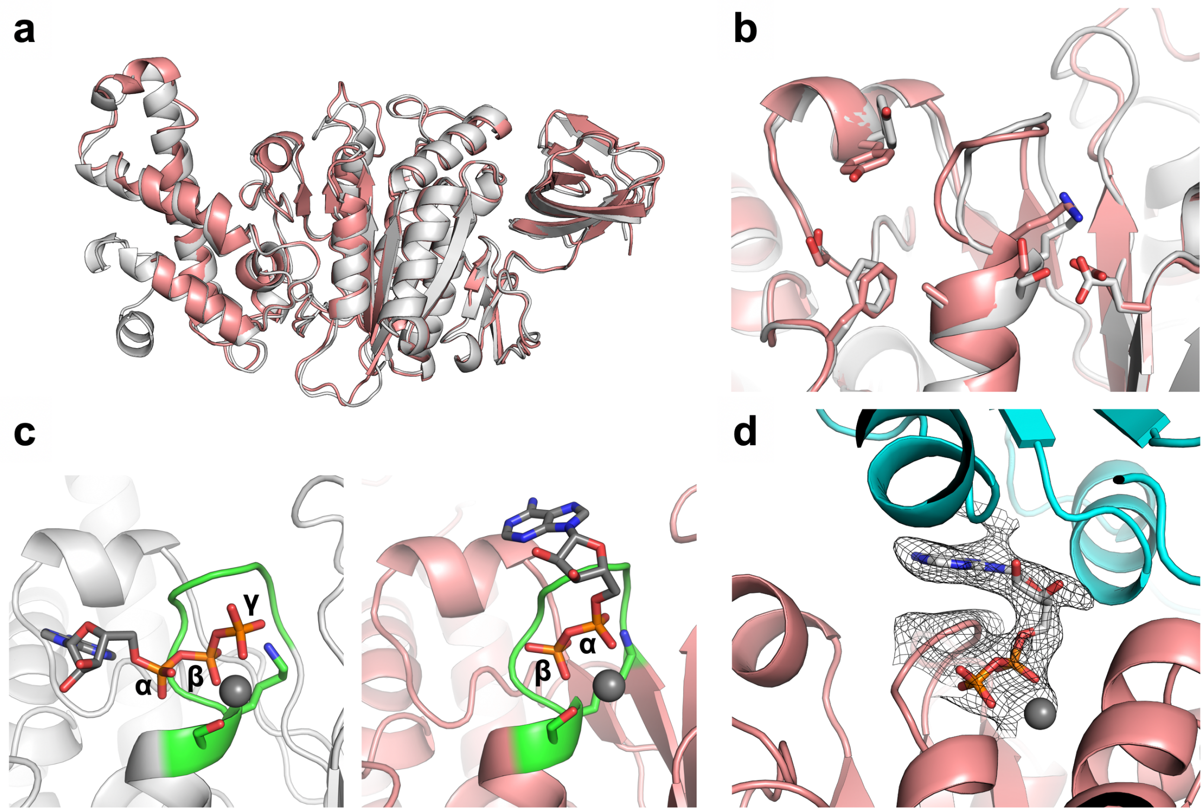


**Supplementary Fig. 9: Structural comparisons of the computational design model with the solved crystal structures.** The computational design model and the solved crystal structures are colored by white and salmon pink, respectively. **a**,**b,** Comparisons with the nucleotide-free crystal structure (the chain D in $A_3(De)_3$_empty) in terms of entire backbones (**a**) and structures at the designed binding site (**b**) presents the overall agreements (entire Cα RMSD value calculated by MICAN[1]: 1.48 Å). **c,** Comparisons of the design model (left) with the nucleotide-bound crystal structure (the chain D in $A_3(De)_3$_$(ADP{\cdot}Pi)_{1cat}(ADP)_{2cat,2non\text{-}cat}$) (right). In the crystal structure, the designed P-loop (green) formed the typical backbone geometries (Supplementary Fig. 3a,b), and a nucleotide (ADP) was found at the designed binding site. However, the binding-mode was different from the design in which the ADP was bound in the opposite direction to the design: the α phosphate was found at the position of the γ phosphate in the design model. **d,** The sugar and base of ADP were found at the interface with A-subunit (cyan) ($2F_o$-$F_c$ map at 1.0σ of ADP and $Mg^{2+}$ molecule is colored by grey). One of the reasons for the unintended ATP binding mode shown in **c** and **d** is likely because we focused on designing the binding site only in the monomer B-subunit without considering the AB complex.

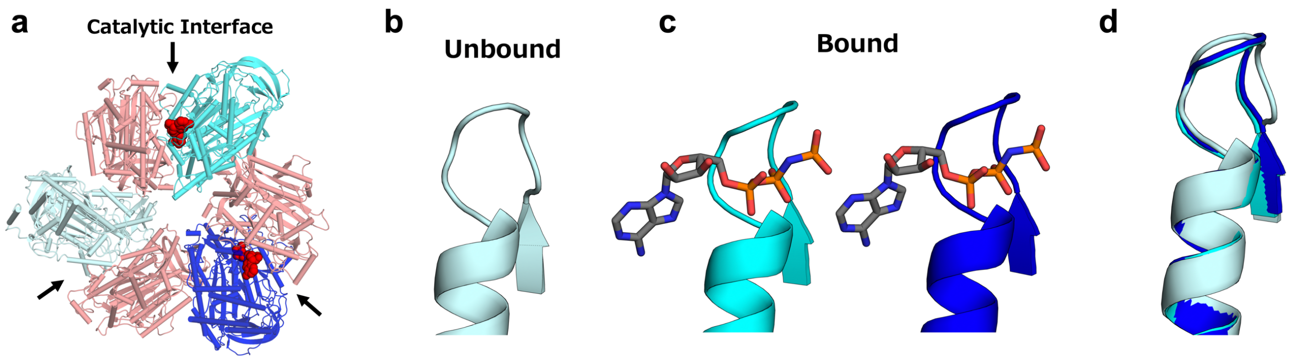


**Supplementary Fig. 10: Bound and unbound forms observed in P-loop motifs in the wild-type structure (PDB: 3VR3). a,** The wild-type $A_3B_3$ complex structure bound to two AMP-PNPs at catalytic interfaces. **b**,**c,** The P-loop motifs present two different types of conformation, depending on the nucleotide binding states at the catalytic interfaces: unbound form (pale cyan) for the nucleotide-free state, and bound form (cyan and blue) for the nucleotide-bound state. **d,** Superposition of the unbound and bound forms.

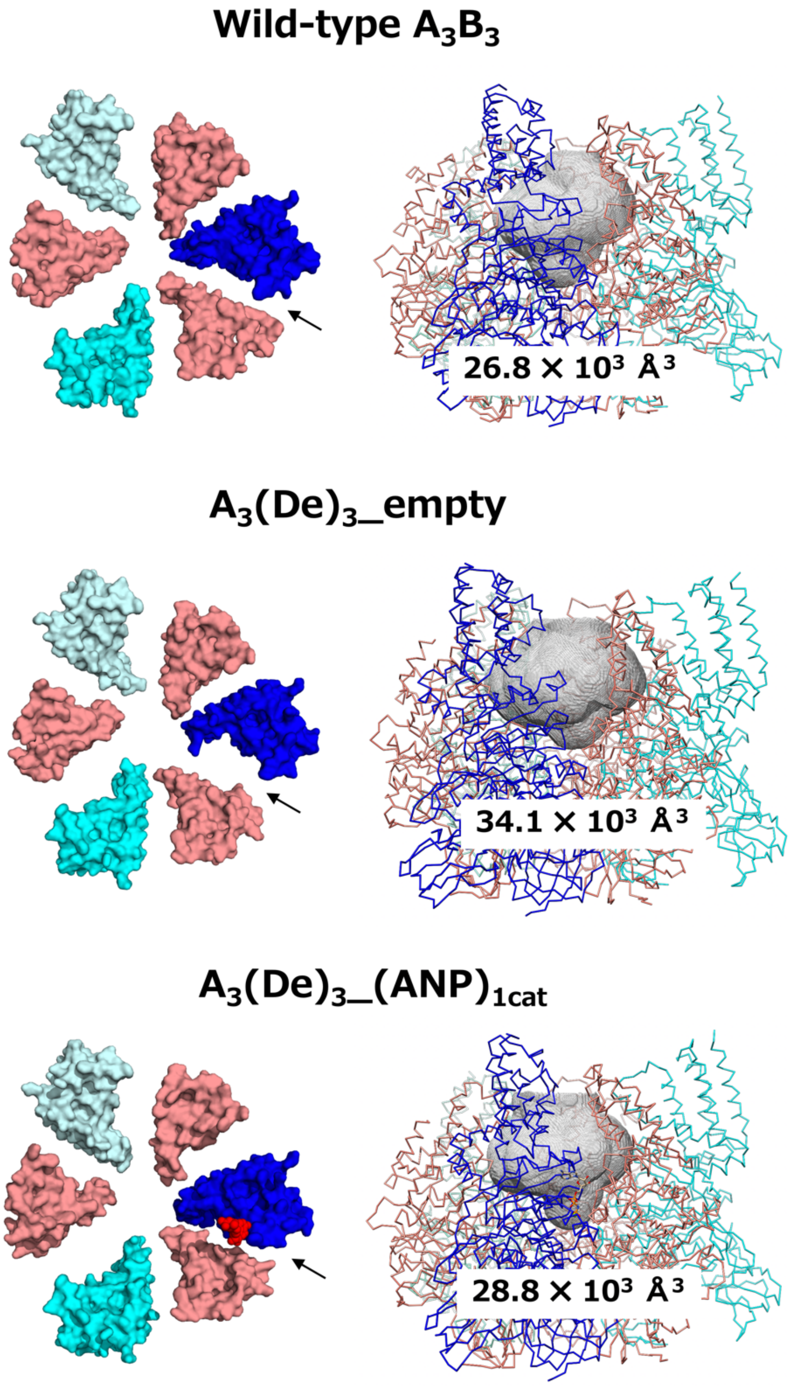


**Supplementary Fig. 11: Design of ATP binding sites in the B-subunit induced conformational changes of the ring complex.** Ring complex conformations of the wild-type $A_3B_3$ complex (top), $A_3(De)_3$_empty (middle) and $A_3(De)_3$_(ANP)$_{1cat}$ (bottom). The C-terminal domains of the A- and B-subunits viewed from the N-terminal β-barrel side, are shown at left. The pore spaces (gray) for central axis are shown in right, which are the views from the arrow of the left figures. The presented pore sizes were calculated using Channel Finder in 3V software[4] with the probe sizes set to 8.0 (small probe) and 16.0 (large probe).

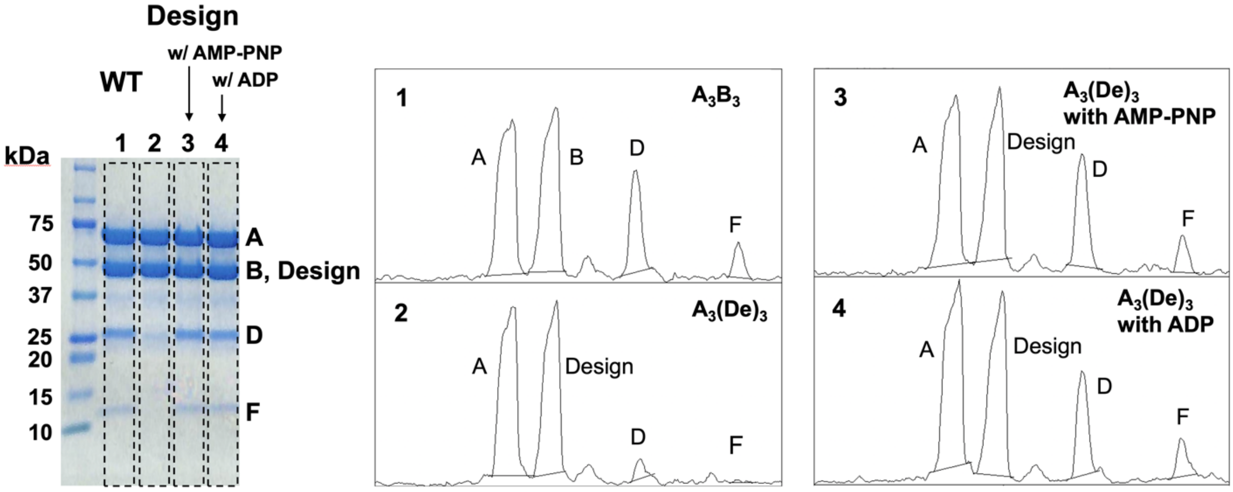


**Supplementary Fig. 12: Complex formation ability of the $A_3B_3$ and $A_3(De)_3$ hexameric ring with the central axis (DF-subcomplex) in the presence or absence of nucleotide.** (Lane 1 and Panel 1) the wild-type $A_3B_3$DF in the absence of nucleotide; (Lane 2 and Panel 2) $A_3(De)_3$DF in the absence of nucleotide; (Lane 3 and Panel 3) $A_3(De)_3$DF in the presence of AMP-PNP; (Lane 4 and Panel 4) $A_3(De)_3$DF in the presence of ADP. The complex reconstitution ratio were evaluated from the SDS-PAGE gel bands (left lanes) by ImageJ[5], in which the pixel intensity of the gel image was converted to the optical density (right panels) by using the following function: uncalibrated OD = $\log_{10}$(255/ pixel value). When the reconstitution rate of the wild-type $A_3B_3$DF is set to 1.0, those of the $A_3(De)_3$DF in the absence of nucleotide, the $A_3(De)_3$DF in the presence of AMP-PNP, and the $A_3(De)_3$DF in the presence of ADP are 0.15, 1.08 and 0.89 respectively.

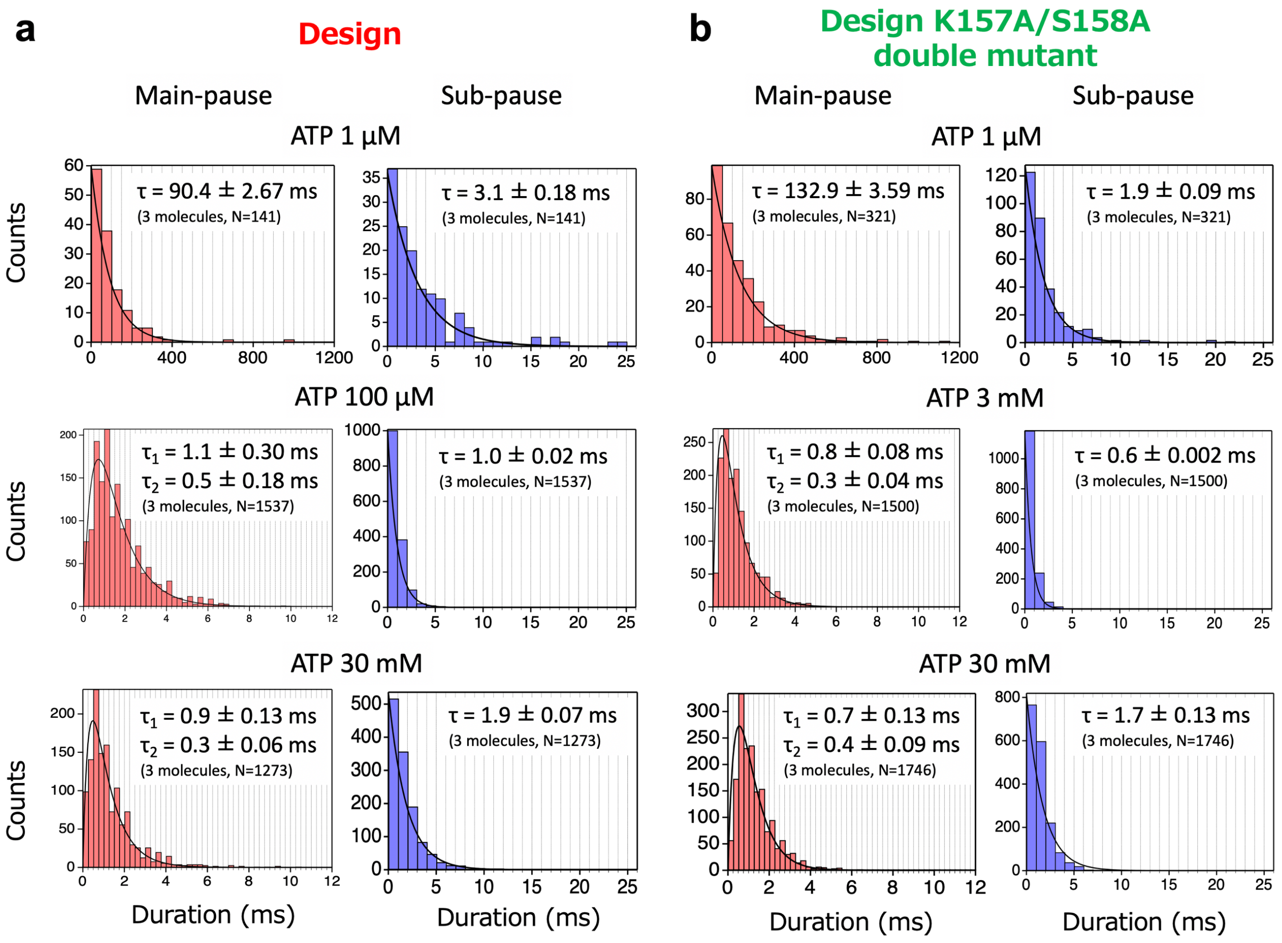


**Supplementary Fig. 13: Duration time distributions with calculated time constants for main- and sub-pauses for the designed $V_1$ and the design double mutant K157A/S158A.** Duration time distribution for main- and sub-pauses for the design (**a**) and the design double mutant K157A/S158A (**b**). For the main-pause at 1 μM [ATP], the distributions were fitted with a single-exponential decay function: constant × exp($-t/\tau$). For the main-pause at 100 μM, 3 mM and 30 mM [ATP], the distributions were fitted with a double-exponential decay functions assuming two consecutive first-order reactions: constant × exp($-t/\tau_1$) - exp($-t/\tau_2$). For the sub-pause at all [ATP]s, the distributions were fitted with a single-exponential decay function: constant × exp($-t/\tau$).

**Supplementary Table 1: Structural comparison of A-subunits between the nucleotide-free $A_3B_3$ complex of the wild-type (3VR2) and the designed $V_1$ ($A_3(De)_3$_empty).**

RMSD values (Å) calculated by MICAN[1] are shown. O, O', C represent conformational states defined in the study by Arai *et al.*[6]; O: open, O': semi open and C: close. The bold numbers indicate the minimal RMSD values for each A-subunit in $A_3(De)_3$_empty. Conformations of all A-subunits in $A_3(De)_3$_empty are close to open (O or O') conformation in the wild-type.

| A subunit Chain | State | $A_3(De)_3$_empty Chain A | $A_3(De)_3$_empty Chain B | $A_3(De)_3$_empty Chain C |
|---|---|---|---|---|
| Wild-type, 3VR2 Chain A | O | **0.35** | 0.70 | 0.94 |
| Wild-type, 3VR2 Chain B | O' | 0.57 | **0.57** | **0.73** |
| Wild-type, 3VR2 Chain C | C | 1.79 | 1.64 | 1.38 |

**Supplementary Table 2: Structural comparison of catalytic interfaces between the nucleotide-free $A_3B_3$ complex of the wild-type (3VR2) and the designed $V_1$ ($A_3(De)_3$_empty).**

RMSD values (Å) calculated by MICAN[1] are shown. Empty, Bindable and Bound represent conformational states of the catalytic interface defined in the study by Arai *et al.*[6]. The bold numbers indicate the minimal RMSD values for each catalytic interface. All catalytic interfaces in $A_3(De)_3$_empty are close to open (Empty or Bindable) conformation in the wild-type.

| Catalytic Interface | State | $A_3(De)_3$_empty Chain AD | $A_3(De)_3$_empty Chain BE | $A_3(De)_3$_empty Chain CF |
|---|---|---|---|---|
| Wild-type, 3VR2 Chain AD | Empty | **1.05** | 2.29 | 2.86 |
| Wild-type, 3VR2 Chain BE | Bindable | 1.36 | **0.95** | **1.73** |
| Wild-type, 3VR2 Chain CF | Bound | 3.52 | 3.39 | 2.42 |

**Supplementary Table 3: Structural comparison of the A-subunits between the nucleotide-free $A_3B_3$ complex of the wild-type (3VR2) and the designed $V_1$ ($A_3(De)_3\_(ANP)_{1cat}$).**

RMSD values (Å) calculated by MICAN[1] are shown. O, O', C represent conformational states defined in the Arai *et al.*[6]; O: open, O': semi open and C: close. The bold numbers indicate the minimal RMSD values for each A-subunit in $A_3(De)_3\_(ANP)_{1cat}$. Conformations of the A-subunits in $A_3(De)_3\_(ANP)_{1cat}$ are similar with those in the wild-type.

| A subunit Chain | State | $A_3(De)_3\_(ANP)_{1cat}$ Chain A | $A_3(De)_3\_(ANP)_{1cat}$ Chain B | $A_3(De)_3\_(ANP)_{1cat}$ Chain C |
|---|---|---|---|---|
| Wild-type, 3VR2 Chain A | O | **0.47** | 0.78 | 2.29 |
| Wild-type, 3VR2 Chain B | O' | 0.54 | **0.64** | 2.25 |
| Wild-type, 3VR2 Chain C | C | 1.80 | 1.79 | **1.24** |

**Supplementary Table 4: Structural comparison of the catalytic interfaces between the nucleotide-free $A_3B_3$ complex of the wild-type (3VR2) and the designed $V_1$ ($A_3(De)_3\_(ANP)_{1cat}$).**

RMSD values (Å) calculated by MICAN[1] are shown. Empty, Bindable and Bound represent conformational states of the catalytic interface defined in the Arai *et al.*[6]. The bold numbers indicate the minimal RMSD values for each catalytic interface in $A_3(De)_3\_(ANP)_{1cat}$. The catalytic interfaces in $A_3(De)_3\_(ANP)_{1cat}$ have similar conformations with those in the wild-type.

| Catalytic Interface | State | $A_3(De)_3\_(ANP)_{1cat}$ Chain AD | $A_3(De)_3\_(ANP)_{1cat}$ Chain BE | $A_3(De)_3\_(ANP)_{1cat}$ Chain CF |
|---|---|---|---|---|
| Wild-type, 3VR2 Chain AD | Empty | **1.19** | 2.71 | 3.82 |
| Wild-type, 3VR2 Chain BE | Bindable | 1.51 | **1.32** | 3.68 |
| Wild-type, 3VR2 Chain CF | Bound | 3.43 | 3.59 | **1.51** |

**Supplementary Table 5: Number of molecules used for measurement of average rotation rates in the single-molecule experiments.**

| [ATP] (μM) | Number of molecules | | |
|---|---|---|---|
| | Designed $V_1$ | Designed $V_1$ K157Q | Designed $V_1$ K157A/T158A |
| 1 | 7 | 3 | 5 |
| 10 | 3 | 3 | 6 |
| 30 | 4 | 4 | 4 |
| 100 | 8 | 4 | 5 |
| 300 | 3 | 3 | 4 |
| 1000 | 4 | 5 | 4 |
| 3000 | 4 | 6 | 9 |
| 10000 | 3 | 5 | 5 |
| 30000 | 5 | 5 | 7 |

**Supplementary Table 6: Comparison of rotation rates estimated from the slope in the time course of rotation and those estimated from the dwell time constants for the main- and sub-pauses.**

| | [ATP] (μM) | Rotation rate estimated from the slope in the time course of rotation | Rotation rate estimated from the dwell time constants |
|---|---|---|---|
| Designed $V_1$ | 1 | 3.0 | 3.6 |
| | 100 | 113.9 | 128.2 |
| | 30000 | 118.0 | 107.5 |
| Designed $V_1$ K157A/S158A | 1 | 2.2 | 2.5 |
| | 3000 | 160.6 | 196.1 |
| | 30000 | 135.7 | 119.0 |

**Supplementary Table 7: Structural comparison of the A-subunits between the nucleotide-free $A_3B_3$ complex of the wild-type (3VR2) and the designed $V_1$ ($A_3(De)_3\_(ADP)_{3cat,1non\text{-}cat}$ and $A_3(De)_3\_(ADP)_{3cat,2non\text{-}cat}$).**

RMSD values calculated by MICAN[1] are shown. O, O', C represent conformational states defined in the Arai *et al.*[6]; O: open, O': semi open and C: close. The bold numbers indicate the RMSD values for the chain with the largest conformational change between $A_3(De)_3\_(ADP)_{3cat,1non\text{-}cat}$ and $A_3(De)_3\_(ADP)_{3cat,2non\text{-}cat}$. One of A-subunits in the design complex changes to more open conformation upon binding a nucleotide at the next designed interface. Note that the RMSD values for chain B in $A_3(De)_3\_(ADP)_{3cat,1non\text{-}cat}$ in parentheses are probably underestimated, since the part of C-terminal helical domain, which mainly moves with the change of conformational state, is deleted in the structural model because of the unclear density.

| A-subunit Chain | State | $A_3(De)_3\_(ADP)_{3cat,1non\text{-}cat}$ | | | $A_3(De)_3\_(ADP)_{3cat,2non\text{-}cat}$ | | |
|---|---|---|---|---|---|---|---|
| | | Chain A | Chain B | **Chain C** | Chain G | Chain H | **Chain I** |
| Wild-type, 3VR2 Chain A | O | 1.31 | (1.27) | **1.80** | 1.14 | 1.53 | **1.49** |
| Wild-type, 3VR2 Chain B | O' | 1.39 | (1.22) | **1.77** | 1.24 | 1.53 | **1.47** |
| Wild-type, 3VR2 Chain C | C | 1.41 | (1.17) | **1.05** | 1.72 | 1.33 | **1.14** |

**Supplementary Table 8: Structural comparison of the catalytic interfaces between the nucleotide-free $A_3B_3$ complex of the wild-type (3VR2) and the designed $V_1$ ($A_3(De)_3\_(ADP)_{3cat,1non\text{-}cat}$ and $A_3(De)_3\_(ADP)_{3cat,2non\text{-}cat}$).**

RMSD values calculated by MICAN[1] are shown. Empty, Bindable and Bound represent conformational states of the catalytic interface defined in the Arai *et al.*[6]. The bold numbers indicate the RMSD values for the catalytic interface with the largest conformational change between $A_3(De)_3\_(ADP)_{3cat,1non\text{-}cat}$ and $A_3(De)_3\_(ADP)_{3cat,2non\text{-}cat}$. One of catalytic interfaces in the design complex changes to more open conformation upon binding a nucleotide at the next designed interface. Note that the RMSD values for chain BE in $A_3(De)_3\_(ADP)_{3cat,1non\text{-}cat}$ in parentheses are probably underestimated, since the part of C-terminal helical domain of chain B is deleted in the structural model because of the unclear density.

| Catalytic Interface | State | $A_3(De)_3\_(ADP)_{3cat,1non\text{-}cat}$ | | | $A_3(De)_3\_(ADP)_{3cat,2non\text{-}cat}$ | | |
|---|---|---|---|---|---|---|---|
| | | Chain AD | Chain BE | **Chain CF** | Chain GJ | Chain HK | **Chain IL** |
| Wild-type, 3VR2 Chain AD | Empty | 2.01 | (2.62) | **3.49** | 2.23 | 2.73 | **2.87** |
| Wild-type, 3VR2 Chain BE | Bindable | 1.98 | (1.48) | **2.82** | 1.42 | 1.84 | **2.05** |
| Wild-type, 3VR2 Chain CF | Bound | 2.61 | (2.55) | **1.62** | 3.21 | 2.50 | **2.09** |

**Supplementary Table 9: Data collection and refinement statistics of crystal structures**

| | $A_3(De)_3\_empty$<br>PDB: 7COP | $A_3(De)_3\_(ANP)_{1cat}$<br>PDB: 7COQ | $A_3(De)_3\_(ADP \cdot Pi)_{1cat}$ $(ADP)_{2cat,2non\text{-}cat}$<br>PDB: 7COR | $A_3(De)_3\_(ADP)_{3cat,1non\text{-}cat}$<br>$A_3(De)_3\_(ADP)_{3cat,2non\text{-}cat}$<br>PDB: 7COP |
|---|---|---|---|---|
| **Data collection** | | | | |
| Space group | $P2_1$ | $P2_1$ | $P2_1$ | $P2_1$ |
| Cell dimensions | | | | |
| *a, b, c* (Å) | 122.45, 122.65, 128.70 | 124.44, 124.49, 131.34 | 119.76, 126.73, 123.81 | 179.44, 125.08, 180.97 |
| α, β, γ (°) | 90.0, 90.7, 90.0 | 90.0, 93.0, 90.0 | 90.0, 94.2, 90.0 | 90.0, 93.8, 90.0 |
| Wavelength | 1.1000 | 1.1000 | 1.1000 | 1.1000 |
| Resolution (Å) | 44.64 – 2.77 (2.82 – 2.77) | 46.34 – 3.44 (3.55 – 3.44) | 44.58 – 2.90 (2.96 – 2.90) | 48.83 – 3.95 (4.04 – 3.95) |
| $R_{merge}$ | 0.09 (0.88) | 0.117 (0.85) | 0.21 (1.86) | 0.28 (2.03) |
| $I/\sigma I$ | 14.2 (2.1) | 11.3 (2.4) | 7.6 (1.0) | 6.8 (1.0) |
| Completeness (%) | 100.0 (100.0) | 99.8 (99.9) | 100.0 (100.0) | 99.9 (99.9) |
| Redundancy | 7.0 (7.3) | 6.9 (6.8) | 6.9 (6.9) | 7.3 (6.4) |
| **Refinement** | | | | |
| Resolution (Å) | 44.64 - 2.77 | 46.34 – 3.44 | 44.62 – 2.90 | 48.83 - 3.95 |
| No. reflections | 96799 | 50840 | 77744 | 70389 |
| $R_{work}/R_{free}$ | 0.22/0.26 | 0.21/0.25 | 0.23/0.29 | 0.24/0.28 |
| No. atoms | | | | |
| Protein | 24061 | 24028 | 24005 | 47582* |
| Ligand/ion | 0 | 32 | 145 | 250 |
| Water | 253 | 0 | 132 | 0 |
| B-factors | | | | |
| Protein | 76.157 | 96.707 | 63.968 | 146.924 |
| Ligand/ion | | 91.085 | 74.488 | 174.963 |
| Water | 56.332 | | 40.158 | |
| R.m.s. deviations | | | | |
| Bond length (Å) | 0.01 | 0.01 | 0.01 | 0.00 |
| Bond angles (°) | 0.92 | 1.00 | 1.59 | 0.60 |

Values in parentheses are for highest-resolution shell.

*Two $A_3(De)_3$ molecules in the asymmetric unit. In the structural models of $A_3(De)_3\_(ADP)_{3cat,1non\text{-}cat}$ and $A_3(De)_3\_(ADP)_{3cat,2non\text{-}cat}$, the density of the C-terminal domain of chain B in $A_3(De)_3\_(ADP)_{3cat,1non\text{-}cat}$ is not clearly observed and the magnesium ions were not placed at the catalytic sites in chain B and L because of the unclear density.

### Supplementary Method

**Rosetta Scripts XML file used for design of ATP binding site:**

```
<ROSETTASCRIPTS>
	<SCOREFXNS>
		<SFXN weights=talaris2014 />
	</SCOREFXNS>
	<FILTERS>
		<Ddg name=ddg_calc scorefxn=SFXN jump=1 threshold=-8 repeats=5 repack=true />
	</FILTERS>
	<TASKOPERATIONS>
		<ReadResfile name=resfile filename="./3VR6_E_Ploop.resfile" />
		<LayerDesign name=layer_all layer=core_boundary_surface core=20 surface_E=70 surface_H=60 pore_radius=2.0 ignore_pikaa_natro=1 />
	</TASKOPERATIONS>
	<MOVERS>
		<AddOrRemoveMatchCsts name=cstadd cst_instruction=add_new/>
		<AddOrRemoveMatchCsts name=cstremove cst_instruction=remove/>
		<EnzRepackMinimize name=min_enz scorefxn_repack=SFXN scorefxn_minimize=SFXN design=1 minimize_bb=1 minimize_sc=1 minimize_rb=1 minimize_lig=1 cycles=20 task_operations=resfile,layer_all />
		<EnzRepackMinimize name=min scorefxn_repack=SFXN scorefxn_minimize=SFXN minimize_sc=1 minimize_rb=0 minimize_lig=1 design=0 repack_only=0 minimize_bb=0 cycles=20 />
		<Idealize name=ideal />
		<PredesignPerturbMover name=pre_min trans_magnitude=0.1 rot_magnitude=2.0 dock_trials=5000 />
	</MOVERS>
	<PROTOCOLS>
		<Add mover_name=ideal />
		<Add mover_name=cstadd/>
		<Add mover_name=pre_min/>
		<Add mover_name=min_enz />
		<Add mover_name=cstremove/>
		<Add mover_name=min/>
		<Add filter_name=ddg_calc />
	</PROTOCOLS>
</ROSETTASCRIPTS>
```
